# Development of a genetically encoded sensor for probing endogenous nociceptin opioid peptide release

**DOI:** 10.1101/2023.05.26.542102

**Authors:** Xuehan Zhou, Carrie Stine, Patricia Oliveira Prada, Debora Fusca, Kevin Assoumou, Jan Dernic, Musadiq A. Bhat, Ananya S. Achanta, Joseph C. Johnson, Amanda Loren Pasqualini, Sanjana Jadhav, Corinna A. Bauder, Lukas Steuernagel, Luca Ravotto, Dietmar Benke, Bruno Weber, Azra Suko, Richard D. Palmiter, Miriam Stoeber, Peter Kloppenburg, Jens C. Brüning, Michael R. Bruchas, Tommaso Patriarchi

## Abstract

Nociceptin/orphanin-FQ (N/OFQ) is a recently appreciated critical opioid peptide with key regulatory functions in several central behavioral processes including motivation, stress, feeding, and sleep. The functional relevance of N/OFQ action in the mammalian brain remains unclear due to a lack of high-resolution approaches to detect this neuropeptide with appropriate spatial and temporal resolution. Here we develop and characterize NOPLight, a genetically encoded sensor that sensitively reports changes in endogenous N/OFQ release. We characterized the affinity, pharmacological profile, spectral properties, kinetics, ligand selectivity, and potential interaction with intracellular signal transducers of NOPLight *in vitro*. Its functionality was established in acute brain slices by exogeneous N/OFQ application and chemogenetic induction of endogenous N/OFQ release from PNOC neurons. *In vivo* studies with fibre photometry enabled direct recording of NOPLight binding to exogenous N/OFQ receptor ligands, as well as detection of endogenous N/OFQ release within the paranigral ventral tegmental area (pnVTA) during natural behaviors and chemogenetic activation of PNOC neurons. In summary, we show here that NOPLight can be used to detect N/OFQ opioid peptide signal dynamics in tissue and freely behaving animals.

## INTRODUCTION

The nociceptin/orphanin-FQ peptide (N/OFQ), along with its cognate receptor (NOPR) encoded by the *Oprl1* gene, represents the most recently discovered opioid peptide/receptor system. ^1,2^ NOPR is a G protein-coupled receptor (GPCR) which shares 60% of sequence similarity with the other members in the opioid family, ^3^ while retaining a unique pharmacological profile. ^4^ Upon occupancy by its endogenous peptide ligand N/OFQ, the receptor activates downstream Gi/Go proteins and induces intracellular signaling that includes the inhibition of cAMP formation, and ultimately reduces neurotransmission via the inhibition of voltage-gated calcium channels and the activation of inwardly-rectifying potassium channels. ^5^

NOPR is abundantly expressed within the central nervous system, ^6,7,8^ in line with the broad range of neural and cognitive functions regulated by this endogenous opioid system. ^9,10^ In particular, NOPR and preproN/OFQ (PNOC)-expressing neurons are highly enriched in the ventral tegmental area (VTA), arcuate nucleus of the hypothalamus (ARC), dorsal striatum, nucleus accumbens (NAc) and medial prefrontal cortex (mPFC). ^7^ Being the major source of dopamine to limbic and forebrain regions, the VTA plays an important part in neural circuits regulating motivation and reward-based learning. ^11,12^ It is well established that VTA dopaminergic projections to the NAc are essential for encoding reward prediction error and adaptive motivated behavior towards both beneficial and aversive stimuli. ^13^ Several studies have shown that the N/OFQ system exerts an important modulatory effect on mesolimbic dopaminergic circuits. For example, intracerebroventricular (ICV) injection of N/OFQ produces a decrease in extracellular dopamine in the NAc ^14^ and exerts an inhibitory constraint on dopamine transmission by either inhibiting tyrosine hydroxylase phosphorylation or dopamine D1 receptor signalling. ^15^ At the behavioral level, N/OFQ has been shown to prevent morphine- and cocaine-induced dopamine increase in the NAc, ^16,17^ and inhibit conditioned place preference to morphine, amphetamine and cocaine, ^18,19^ while disruption of the N/OFQ system is associated with motivated responding disorder. ^20^ ICV administration of N/OFQ was reported to potently block reward-associated cues but showed no effect on aversion associated cues. ^21^

In a recent study, we identified a subgroup of PNOC-enriched neurons located in the paranigral VTA which, when activated, caused avoidance behavior and decreased motivation for reward. ^22^ Additionally, a separate population of PNOC neurons located in the ARC has emerged as an important neuronal population involved in regulating feeding behavior. These GABA-expressing neurons are activated after three days of a high-palatable, energy-dense diet and have been found to play a crucial role in feeding control. In particular, optogenetic stimulation of PNOC-expressing neurons in the ARC induces feeding, while selective ablation of these neurons decreases food intake and prevents obesity. ^23^ Apart from its roles in the VTA-NAc and ARC circuits, N/OFQ can also inhibit mPFC-projecting VTA neurons ^24^ and a reduction in mPFC N/OFQ level was reported in rodents that underwent conditioned opioid withdrawal. ^25^

Of note, past studies on N/OFQ signalling have generated some contradictory results. ^26^ Activation of the NOPR with a selective agonist was reported to reduce alcohol drinking and seeking behavior, ^27,28^ while a selective NOPR antagonist, LY2940094, was reported to have the same effect. ^29^ In anxiety-related behaviors, it has been reported that central injection of a NOPR agonist induces anxiogenic effect, but anxiolytic effects of NOPR agonists had also been reported. ^30,31,32^ These observations can be interpreted in different ways. The dynamics of NOPR desensitization after application of agonists or antagonists, for example, could contribute to these contradictory results. Another possible explanation could be the competition of different local neural circuits simultaneously recruited by the N/OFQ system, as most of the studies mentioned before do not have fine spatial control over the application of drugs nor the resolution to isolate endogenous release dynamics of the peptide. Overall, the exact mechanism of the NOPR-N/OFQ system and its impact on different neural circuits are, at best, only partially understood.

A major factor hindering a clearer understanding of N/OFQ regulation of neural circuits, or any neuropeptide signalling system, is the limitations imposed by current tools and techniques used to detect the release of neuropeptides in living systems. Conventional techniques such as microdialysis and mass spectrometry-coupled high-performance liquid chromatography can successfully detect picomolar level of neuropeptides in extracellular fluid, ^33^ yet the spatial and temporal resolution of these techniques is limited to single point measurements and long timescales on the order of minutes. ^34^ This temporal and spatial resolution is low for decoding the neuronal mechanisms of dynamic peptide action *in vivo*, particularly alongside other neurophysiological methods. New approaches to probe the nervous system using fluorescent sensors have started to gain traction across the field. ^35,36,37,38,39^ Combined with rapidly developing fluorescent recording and imaging techniques, these sensor-based approaches are uniquely suited for *in vivo* observations with finer spatiotemporal resolution than was previously possible. ^35,40,41^

Here we report the development and characterization of NOP-Light, a novel genetically encoded opioid peptide sensor that provides a specific and sensitive fluorescence readout of endogenous N/OFQ dynamics with unprecedented temporal resolution *ex vivo* and *in vivo*. Using the sensor, we could detect ligand binding by systemically administered NOPR agonists and antagonists within the central nervous system. We also measured both chemogenetically-evoked and behaviorally induced dynamics of endogenous N/OFQ in freely moving mice. Thus, NOPLight extends the neuropeptide molecular toolbox necessary to investigate the physiology of neuropeptides and in particular this important endogenous opioid system with high resolution.

## RESULTS

### Development of a genetically encoded N/OFQ opioid peptide (N/OFQ) sensor

To develop a fluorescent sensor for N/OFQ, we started by designing a prototype sensor based on the human NOPR which has 93-94% sequence identity to the mouse and rat receptors. We replaced the third intracellular loop (ICL3) of human NOPR with a circularly-permuted green fluorescent protein (cpGFP) module that was previously optimized during the development of the dLight1 family of dopamine sensors41 (**Fig. 1a, Supplementary Fig. 1a**). This initial construct exhibited poor membrane expression and no fluorescent response to N/OFQ (**Supplementary Fig. 1b**). Given the pivotal role of the GPCR C-terminus in trafficking, ^43^ we reasoned that replacement of the NOPR C-terminus with that of another opioid receptor may facilitate membrane targeting of the sensor. Based on our prior experience, ^41^ we chose to use the C-terminus from the kappa-type opioid receptor. The resulting chimeric receptor showed improved expression at cell surface, but still exhibited only a small response to N/OFQ (**Supplementary Fig. 1b**).

**FIGURE 1.**
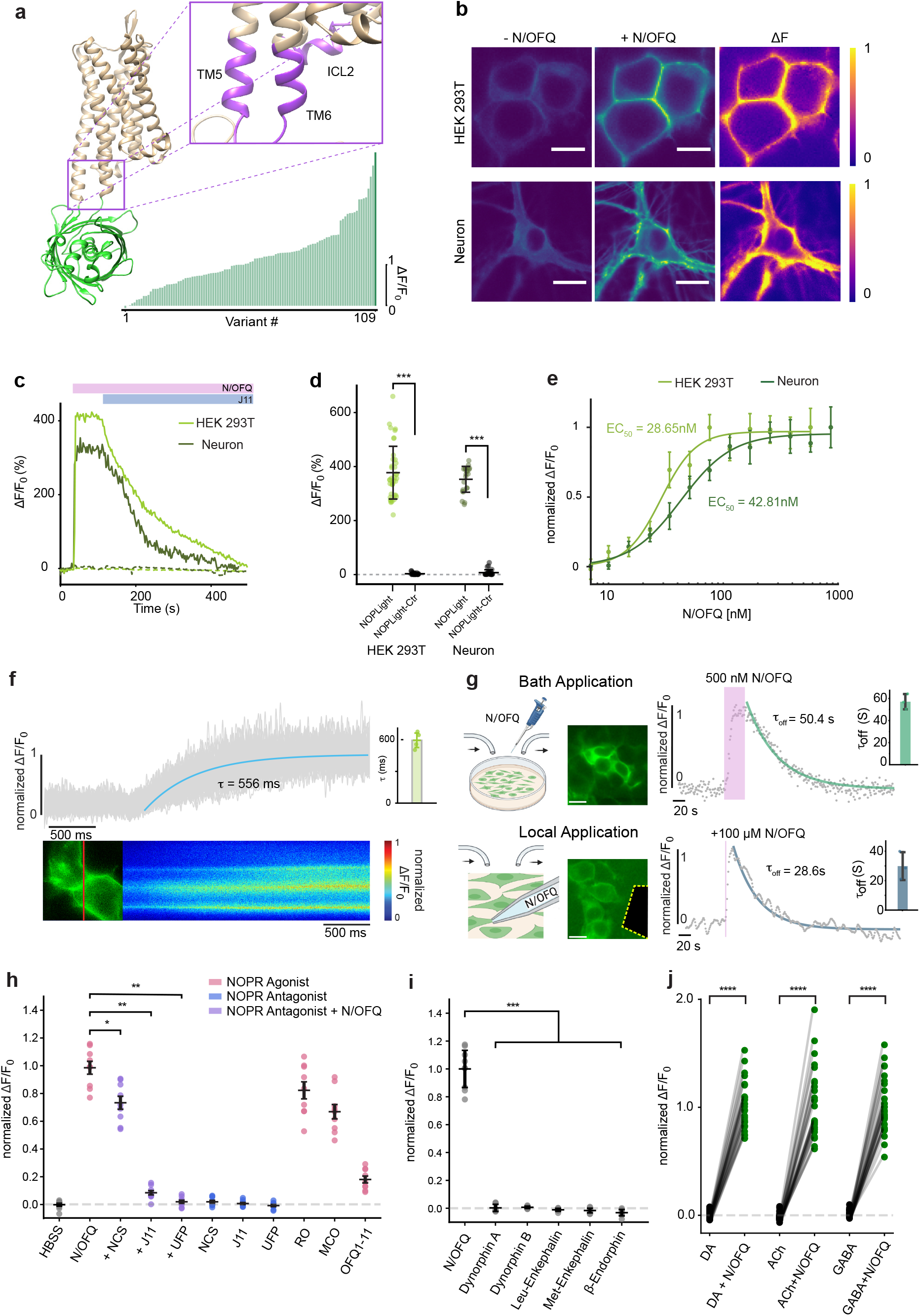
Development and *in vitro* characterization of NOPLight. **a**. Upper left: RoseTTAFold ^42^ predicted structural model of NOPLight. Upper right: zoom in of receptor and cpGFP linker region. Residues of site-directed mutagenesis highlighted in purple. Lower panel: Summary of maximal ΔF/F_0_ in response to 10 μM N/OFQ of HEK293T cells expressing each of the 109 variants screened in this study. (n ≥ 3 cells for each variant. Dark green bar: NOPLight). **b**. Representative images of HEK 293T cells (top; scale bars, 10 μm) and neurons (bottom; scale bars, 20 μm) expressing NOPLight before (left) and after (middle) application f 1 μM N/OFQ. Corresponding normalized pixelwise ΔF shown on the right. **c**. Representative fluorescent-fold change (ΔF/F_0_) of HEK 293T cell (light green traces) and neurons (dark green traces) expressing NOPLight (solid line) or NOPLight-ctr (dotted line) in response to 1 μM N/OFQ, followed by competition of 10 μM J-113397, a NOPR antagonist. **d**. Quantification of maximal ΔF/F_0_ of NOPLight (green) or NOPLight-ctr (gray) expressing HEK293T cells and neurons in response to 1 μM N/OFQ. (n = 38, 30, 22, 27 cells from >3 independent experiments, left to right respectively. Data shown as mean ± s.t.d.) ***P<0.0001. P = 1.003 x 10-12 and 1.262 x 10-9 (One-sided Mann-whitney U test) for the response of NOPLight compared to NOPLight-ctr in HEK 293T cells and neurons. **e**. Normalized maximal ΔF/F_0_ response of HEK 293T cells (light green) and neurons (dark green) expressing NOPLight to different concentrations of N/OFQ (Data shown as mean ± s.d.) and respective dose response curve fitted with a three parameter Hill equation. n = 3 independent experiments with > 5 cells each. **f**. Time plot of normalized single NOPLight pixel ΔF/F_0_ (gray) from a representative line-scan (upper right). Pixel average ΔF/F_0_ were fitted with a mono-exponential function (blue trace) and the deduced time constant (τ). Corresponding cell image (red: line scanned) and time profile of all pixels on the line scanned are shown directly under the time plot. Upper right inset: quantification of time constant (τ) from four independent experiments. **g**. Left: schematic representation of the experimental set-ups; middle: Representative images of HEK 293T cells (scale bars, 20 μm), right: time plot of of normalized single NOPLight ΔF/F_0_ from a representative experiment (gray) with fitted mono-exponential decay function (green and blue) with the deduced time constant (τ_off_). Shaded pink bar represents the application of N/OFQ. Upper inset: quantification of time constant (τ_off_) from three independent experiments. **h**. Maximal ΔF/F_0_ response in NOPLight expressing HEK293T cells to the application of different drug(s) (NCS: Nocistatin; J11: J-113397; UFP: UFP-101; RO: Ro 64-6198; MCO: MCOPPB; OFQ1-11: Orphanin FQ (1-11)). Response to N/OFQ in the presence of each antagonist was compared to where only N/OFQ was applied by a pairwise Mann-Whitney rank test with post hoc Bonferroni correction (*P<0.01, **P<0.005.P = 0.024, Nocistatin; 0.0012, J-113397; 0.0012, UFP 101;) **i**. Maximal ΔF/F_0_ response in NOPLight expressing HEK293T cells to the application of endogenous opioid ligands (1 μM). Response to each ligand was compared to the response to N/OFQ by a pairwise Mann-Whitney rank test with post hoc Bonferroni correction (**P<0.005, P = 0.003, Dynorphin A; 0.003, Dynorphin B; 0.003, Leu-Enkephalin ; 0.003, Met-Enkephalin ; 0.003, β-endorphin.) **j**. Maximal ΔF/F_0_ response in NOPLight expressing HEK293T cells to the application of fast neurotransmitters (DA: dopamine, ACh: acetylcholine, GABA: gamma-Aminobutyric acid at 1 mM) normalized to its maximal ΔF/F_0_ response to N/OFQ (1 μM). n = 29-30 cells from 3 independent experiments. ****P<0.0001.

We then aimed to improve the dynamic range of the sensor through mutagenesis efforts. First, we elongated the N-terminal cpGFP linker with additional amino acids originating from dLight141. This led to the identification of a variant with a fluorescent response (ΔF/F_0_) of approximately 100% (**Supplementary Fig. 1c-d**). Prior work demonstrated that sequence variations in the second intracellular loop (ICL2) of the sensor can effectively be used to modulate sensor response. ^41,44^

Thus, as a next step, we performed targeted mutagenesis focusing on the ICL2 of the sensor. Through these efforts we identified a beneficial mutation (I15634.51K, **Supplementary Fig. 1e**) that was then carried forward onto the next rounds of screening, which focused on receptor and cpGFP residues around the insertion site of the fluorescent protein between transmembrane helixes 5/6 (TM5/TM6) (**Supplementary Fig. 1e**). The final variant, which we named NOPLight, had a ΔF/F_0_ of 388% in transfected HEK293T cells and a similar performance in transduced neuron culture (ΔF/F_0_ = 378%) upon activation by the high affinity, full agonist N/OFQ (**Fig. 1b**). Furthermore, the evoked fluorescence signal could be reversed to baseline levels using the selective and competitive small molecule NOPR antagonist J-113397 (**Fig. 1c**).

To aid the subsequent characterization experiments we also developed a control sensor, NOPLight-ctr, by mutating into alanine two key residues (D1102.63, D1303.32) located in the binding pocket of NOPR4 (**Supplementary Fig. 2a, Supplementary Data S1**). The control sensor was well expressed on the surface of HEK293T cells and neurons but showed negligible fluorescent response to N/OFQ (**Fig. 1c-d, Supplementary Fig. 2a**) and a panel of other endogenous opioids and fast neurotransmitters, including dopamine, acetylcholine and GABA. (**Supplementary Fig. 2b**).

**FIGURE 2.**
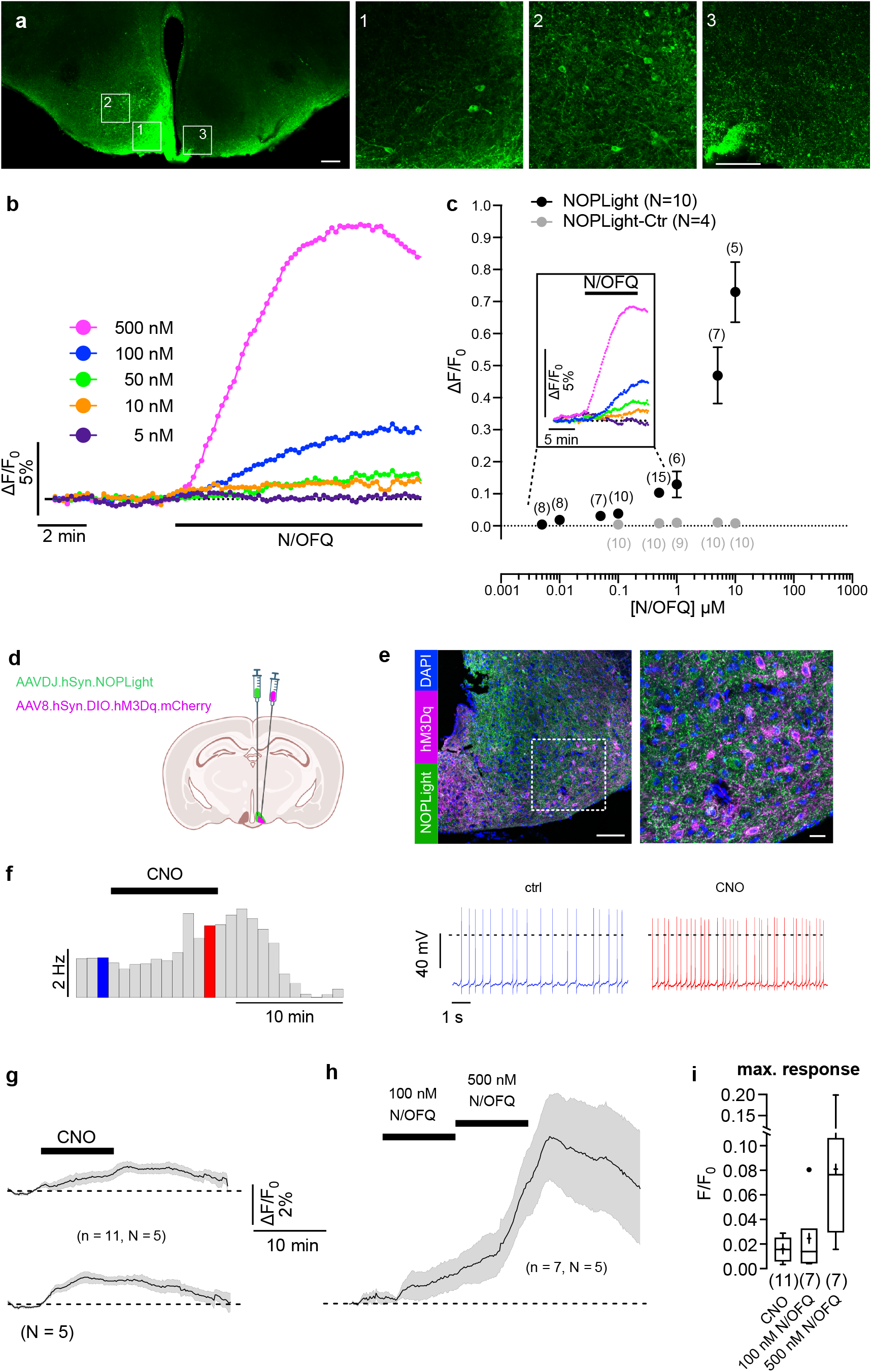
*Ex vivo* characterization of NOPLight. **a**. Expression of NOPLight in the ARC. Scale bar, 200 μm. Insets, scale bars, 50 μm. **b-f**. Change in NOPLight fluorescence measured in ARC neurons in response to bath-applied N/OFQ or release of chemogenetically-activated PNOC neurons. The fluorescence was detected from 0.15– 0.2 mm^2^ ROIs in the ARC. Each experiment was performed in a different brain slices. **b**. NOPLight responses of a single ROI to increasing N/OFQ concentrations. **c**. Concentration-response relation showing the N/OFQ (black trace) effect on N/OFQ fluorescence of NOPLight-expressing cells in the ARC. The grey trace shows the concentration-response relation for the ligand insensitive mutant sensor (NOPLight-ctr, grey trace). Data are shown as mean ± SEM. Inset, mean fluorescence increase in response to different concentrations of N/OFQ. Color code as in **b**. ARC, arcuate nucleus of the hypothalamus. **d**. Experimental schematic for chemogenetic experiments in brain slices. **e**. Representative immunohistochemical image showing DREADD (hM3Dq) expression in PNOC neurons of the ARC as well as pan-neuronal expression of NOPLight. Scale bar, 100 μm. Magnification of inset is shown on right. Scale bar, 20 μm. **f**. Perforated patch clamp recordings from a hM3Dq-expressing PNOC neuron showing the effect of 10 min CNO application (3 μM) on action potential frequency. Left, rate histogram (bin width 60s). Right, representative sections of the original recording corresponding to the times indicated by blue and red color code. **g-i**. Changes (mean ± SEM) in NOPLight fluorescence measured in the ARC in response to activation of hM3Dq-expressing PNOC neurons by 10 min bath-application of 3 μM clozapine N-oxide (CNO) (**g**) and 100 nM and 500 nM N/OFQ (**h**). **g**. The upper and lower panel show the same data. The traces in the upper panel are aligned to the CNO application, the traces in the lower panel are aligned to the response onset. The recordings in **g** and **h** were performed from the same brain slices. The n-values in **h** are lower than in **g** because some recordings were terminated for technical reasons. Bars indicate the application of CNO and N/OFQ, respectively. Scale bars apply to **g** and **h. i**. Box plots showing the maximal fluorescence changes upon applications of CNO and N/OFQ, respectively. The numbers in brackets represents the numbers of experiments (brain slices). N-values indicate the number of animals.

### *In vitro* characterization of NOPLight

To better examine the pharmacological and kinetic properties of the sensor, we first characterized the apparent ligand affinity of NOP-Light *in vitro*, using NOPLight-expressing HEK293T cells and cultured neurons. In HEK293T cells, the endogenous ligand N/OFQ elicited a fluorescent response of NOPLight at a half maximal effective concentration (EC_50_) of 28.65 ± 5.1 nM (pEC_50_ = 7.54), whereas in cultured neurons it showed an EC_50_ of 42.81 ± 5.4 nM (pEC_50_ = 7.37) (**Fig. 1e**), approximately one order of magnitude lower than the reported potency of N/OFQ to the wild type NOPR in the central nervous system. ^45^ To determine the activation kinetics of NOPLight, we measured the activation of NOPLight upon direct bath-application of N/OFQ using high-speed line-scan confocal imaging. Mono-exponential fitting of NOPLight fluorescent response indicated a subsecond time constant of signal activation at the sensor (τ_ON_ = 595 ± 69 ms; **Fig. 11f**). To determine the off kinetics of the sensor, we performed experiments on cells under constant bath-perfusion. Depending on the experimental set-up, the fluorescence response of NOPLight elicited by N/OFQ application returned to baseline with a τ_off_ of 57.1 ± 6.9 s for bath application and 29.9 ± 9.5 s for localized puff application of the ligand (**Fig. 1g**).

We characterized the pharmacological profile of NOPLight *in vitro*. We tested the response of NOPLight-expressing HEK293T cells to a panel of small-molecule and peptide ligands that are known NOPR agonists or antagonists (**Fig. 1h-j**). Of the antagonist compounds tested, the antagonist peptide UFP-101 and the small molecule compound J-113397 produced robust competitive antagonism, fully reversing the activation of NOPLight at the concentrations used (1 μM), while nocistatin elicited a smaller decrease of the signal induced by N/OFQ. Importantly, none of the antagonistic ligands elicited a noticable fluorescent response when applied alone to sensor-expressing cells. On the other hand, we could clearly detect positive fluorescent responses of NOP-Light to several types of selective NOPR agonist compounds. In particular, the full agonist Ro-64 elicited the largest fluorescent response in this assay (ΔF/F_0_ = 323%) and produced a response of similarly large magnitude in NOPLight-expressing primary neuronal cultures (ΔF/F_0_ = 221%) (**Supplementary Fig. 2c-f**). Interestingly, all of the agonist compounds tested induced an overall smaller fluorescence response than N/OFQ itself, when applied at the same concentration (**Fig. 1h**).

We then characterized the spectral properties of NOPLight and NOPLight-ctr. Under both one-photon and two-photon illumination, NOPLight exhibited similar spectral characteristics as other GPCR-based sensors. ^41,46^ The sensor had an isosbestic point at around 440 nm, and peak performance, measured as the ratio between N/OFQ bound versus unbound state, at 472 nm and 920/990 nm, respectively (**Supplementary Fig. 3a-b**). When tested with another NOPR agonist, RO 64-6198, the sensor exhibited similar one-photon spectral properties, whereas the control sensor showed little difference in excitation and emission in the presence or absence of the ligands (**Supplementary Fig. 3a**).

**FIGURE 3.**
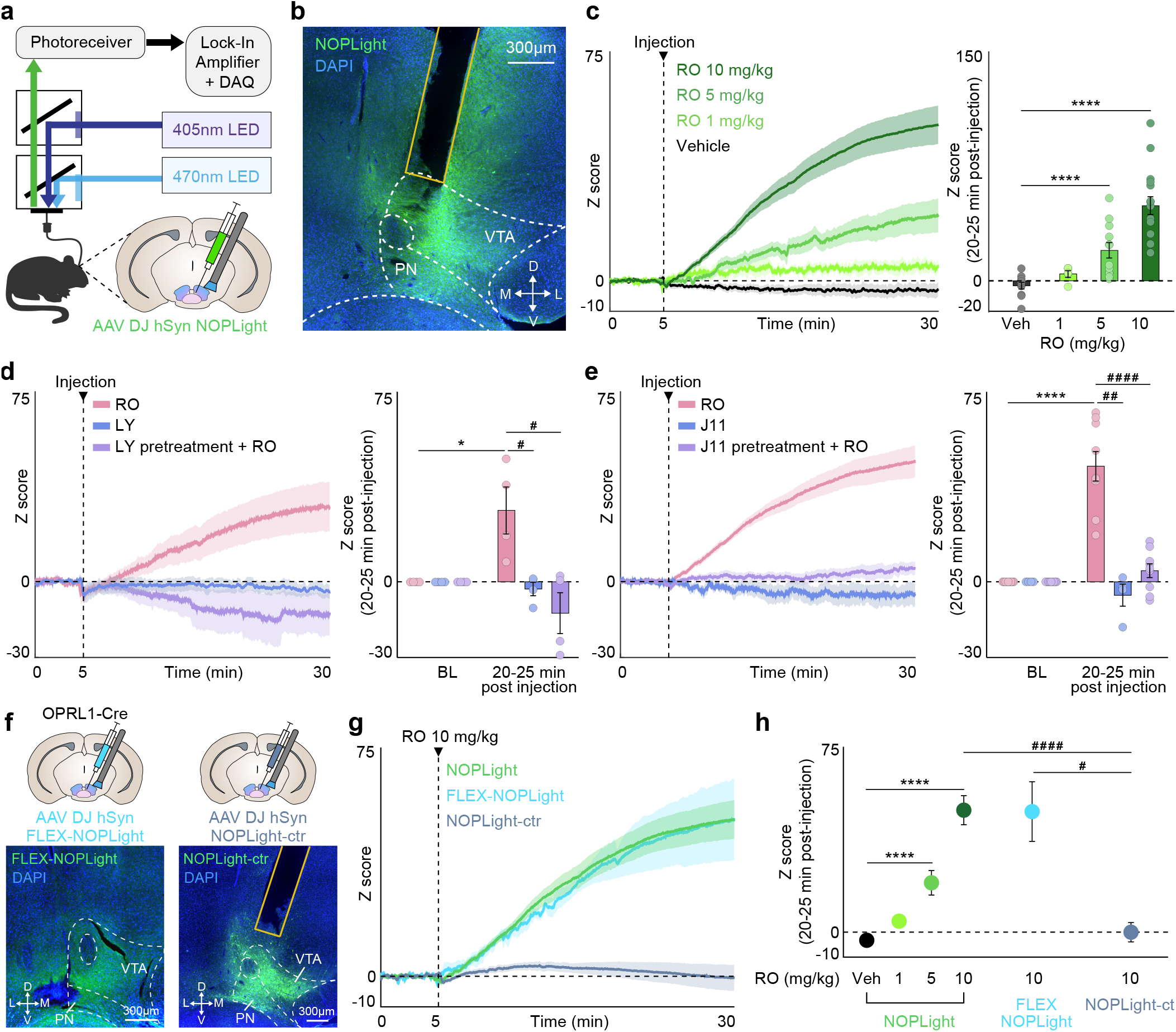
Characterization of NOPLight response *in vivo* to pharmacological agonism and antagonism. **a**. Schematic of fibre photometry setup. Coronal brain cartoon of viral injection of NOPLight and fibre implant in the VTA. **b**. Representative image showing expression of DAPI (blue) and NOPLight (green) with fibre placement in VTA. **c**. Left: Averaged traces of NOPLight fluorescence after systemic (i.p.) injection of vehicle (black) or 1, 5, or 10 mg/kg of selective NOPR agonist Ro 64-6198. Right: Mean NOPLight fluorescence 20-25 min after Ro 64-6198 injection increases dose-dependently (two-tailed Mann-Whitney test, ****p<0.0001, n = 5-16 mice). Data represented as mean ± SEM. **d**. Left: Averaged traces of NOPLight fluorescence after systemic injection of Ro 64-6198 (i.p.) and/or 10 mg/kg of selective NOPR antagonist LY2940094 (o.g.). LY2940094 was administered to the LY + RO group 30 min prior to photometry recording. Right: A significant increase in NOPLight fluorescence 20-25 minutes following injection of Ro 64-6198 (two-tailed Mann-Whitney test, *p<0.05, n = 4 mice) is blocked by NOPR antagonist pre-treatment (two-tailed Mann-Whitney test, # p<0.05, n = 4 mice). Data represented as mean ± SEM. **e**. Left: Averaged traces of NOPLight fluorescence after systemic (i.p.) injection of Ro 64-6198 and/or 10 mg/kg of selective NOPR antagonist J-113397. J-113397 was administered to the J11 + RO group 30 min prior to photometry recording. Right: NOPR antagonist pre-treatment blocks Ro 64-6198 induced increases in NOPLight fluorescence (two-tailed Mann-Whitney test, ## p<0.01, #### p<0.0001, **** p<0.0001, n = 4-9 mice). Data represented as mean ± SEM **f**. Top: Coronal brain cartoon of viral injection of FLEX-NOPLight (left) or NOPLight-ctr (right) and fibre implant in the VTA. Bottom: Representative image showing expression of DAPI (blue) and FLEX-NOPLight (left, green) or NOPLight-ctr (right, green) with fibre placement in VTA. **g**. Averaged traces of NOPLight (green), FLEX-NOPLight (blue) or NOPLight-ctr (gray) fluorescence after systemic (i.p.) injection of 10 mg/kg Ro 64-6198 (n = 3-16 mice). Data represented as mean ± SEM. **h**. Mean fluorescence of each NOPLight variant 20-25 min after systemic injection of Ro 64-6198 (two-tailed Mann-Whitney test, # p<0.05, #### p<0.0001, n = 3-16 mice). Data represented as mean ± SEM.

To evaluate the effect of p.H. change on the sensor, we measured the fluorescence intensity and response of NOPLight-expressing cells to N/OFQ when the cells were exposed to buffer solutions at set p.H. values (6 – 8). Under these conditions, there was no significant difference in sensor response across the tested p.H. range in comparison to neutral p.H. (P = 0. 942, 0.358, 0.883 and 0.289 for p.H. 6, 6.5, 7.5, 8, versus p.H. 7, respectively; one-way ANOVA with Tukey Kramer post-hoc test, **Supplementary Fig. 4a, b**). We also acquired one-photon excitation and emission spectra of NOPLight under similar conditions. Within the tested p.H. range, the isosbestic point of NOPLight consistently fell within the range of 405 - 435 nm. Furthermore, the ratio between N/OFQ-bound versus unbound states also remained overall unaffected by the change in extracellular pH (**Supplementary Fig. 4c**).

**FIGURE 4.**
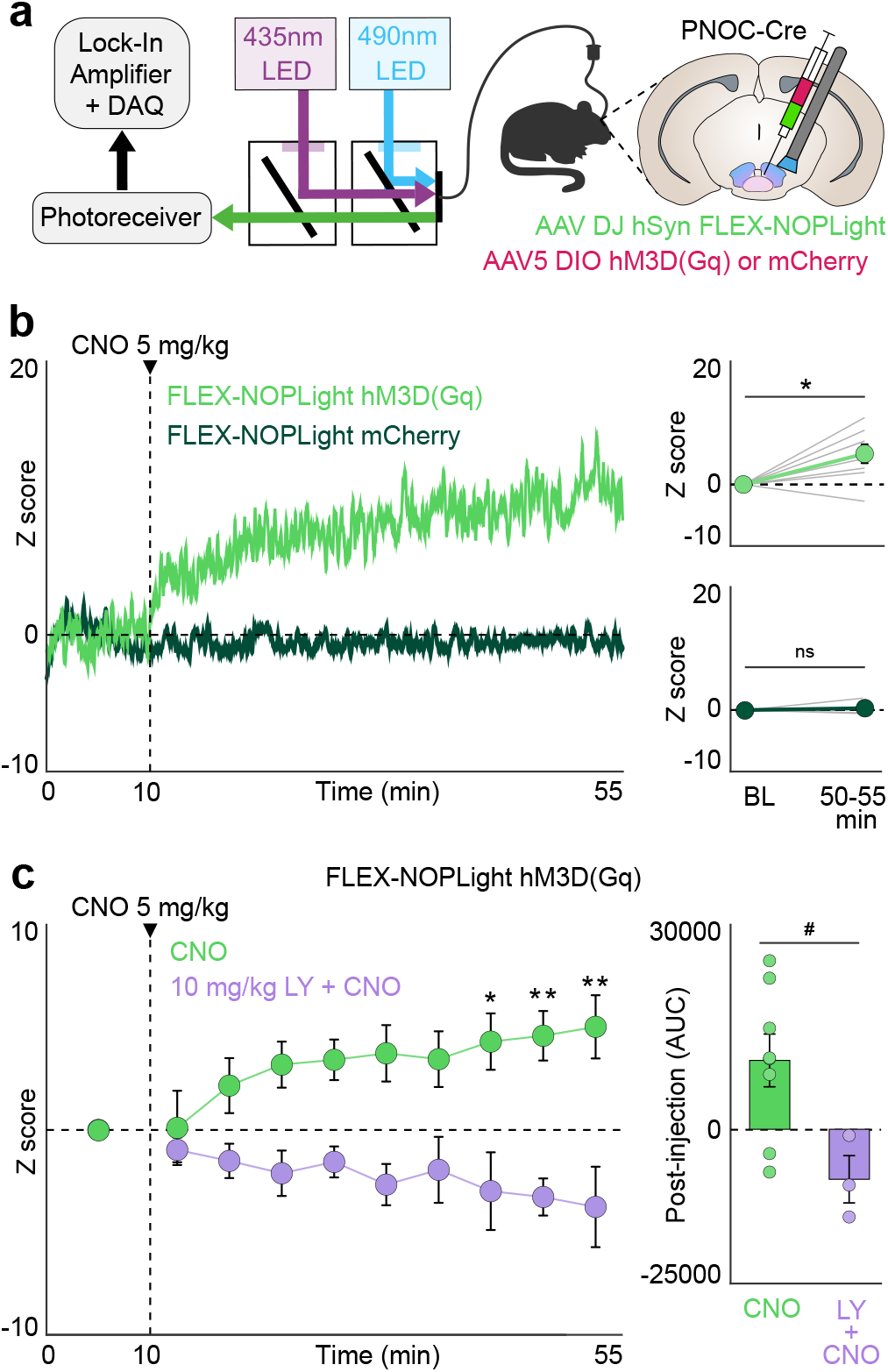
NOPLight detects chemogenetically evoked endogenous N/OFQ release *in vivo*. **a**. Schematic of fibre photometry setup. Coronal brain cartoon of viral co-injection of FLEX-NOPLight with either DIO-hM3D(Gq) or mCherry and fibre implant in the VTA of PNOC Cre mice. **b**. Left: Representative traces of FLEX-NOPLight fluorescence after systemic (i.p.) injection of 5 mg/kg CNO in hM3D(Gq) (green) or control (red) animals. Right: Mean FLEX-NOPLight fluorescence 40-45 min after CNO injection is significantly elevated relative to pre-injection baseline period (BL) in hM3D(Gq) but not control mice (two-tailed Wilcoxon test, *p<0.05, n = 8 mice, hM3D(Gq); 3 mice, control). Data represented as mean ± SEM. **c**. Left: FLEX-NOPLight fluorescence averaged before injection (i.p.) of 5 mg/kg CNO (0-10 min), and in 5 min bins following the injection.10 mg/kg of selective NOPR antagonist LY2940094 was administered (o.g.) to the LY + CNO group 30 min prior to photometry recording (two-way repeated-measures ANOVA with Bonferroni’s post hoc test, *p<0.05, **p<0.01, n = 3-8 mice). Right: NOPR antagonist pre-treatment prevents CNO induced increases in FLEX-NOPLight fluorescence (two-tailed Mann-Whitney test, # p<0.05, n = 3-8 mice). Data represented as mean ± SEM.

Lastly, we evaluated whether changes in neural activity could cause alterations in the observed fluorescence of NOPLight. To do this we co-expressed NOPLight with the red genetically encoded calcium indicator JRCaMP1b ^47^ in primary cultured neurons via viral transduction, followed by simultaneous multiplexed imaging of NOPLight fluorescence and neuronal calcium activity in the absence and presence of bath-applied glutamate at a concentration known to evoke neuronal activity (5 μM). ^48^ Overall, we did not observe noticeable changes in the fluorescence of NOPLight during periods of evoked neuronal activity (**Supplementary Figure 5**), suggesting that potential alterations in the intracellular environment occurring during neuronal activity are not likely to influence the fluorescent responses of NOPLight.

**FIGURE 5.**
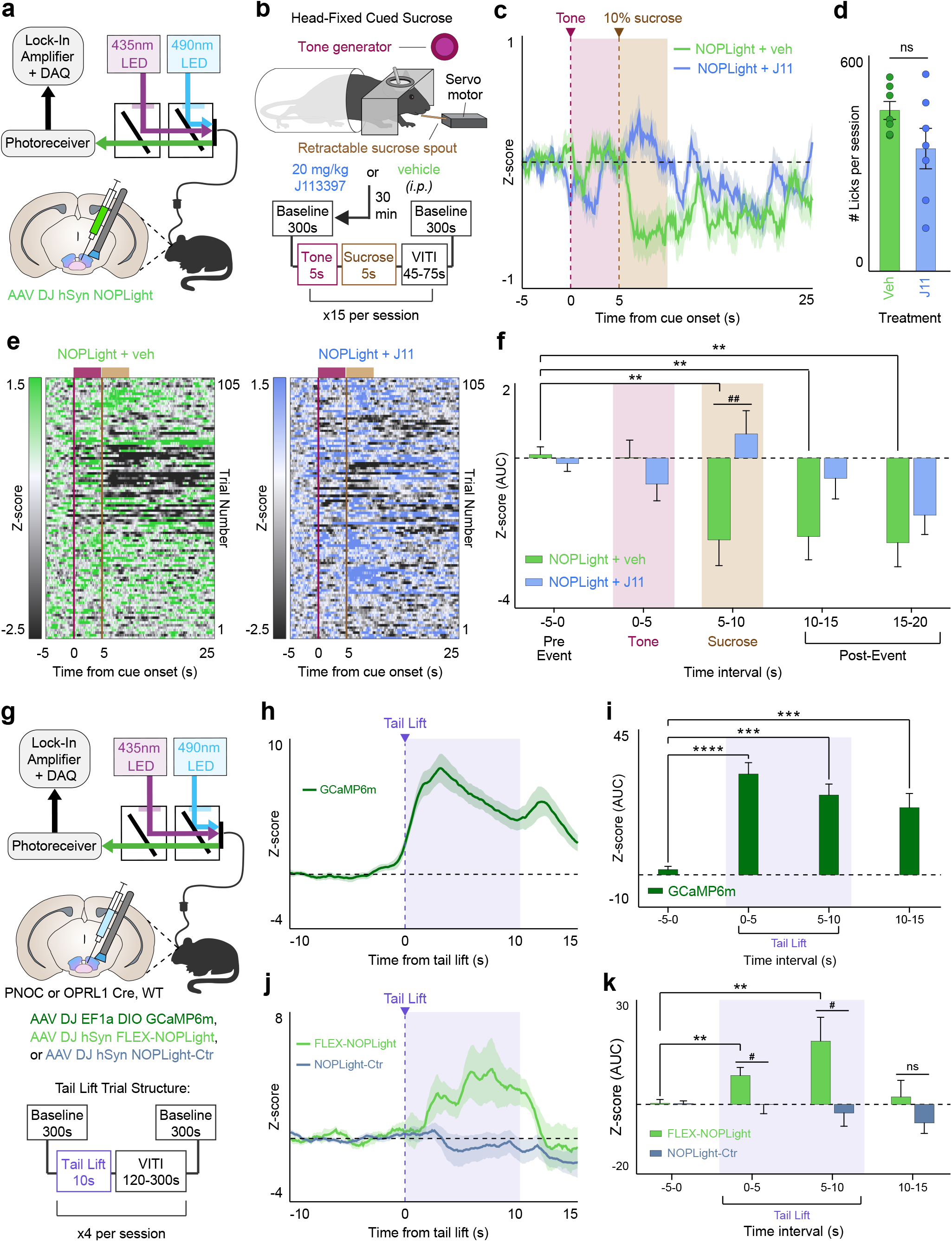
NOPLight *in vivo* reports bidirectional changes in endogenous N/OFQ released during consummatory and aversive behaviors. **a**. Schematic of fibre photometry setup. Coronal brain cartoon of viral injection of NOPLight and fibre implant in the VTA of WT mice (n = 7). **b**. Top: Cartoon depicting setup used for the head-fixed cued sucrose recordings. Bottom: Trial structure for head-fixed cued sucrose sessions. Mice were pre-treated with 20 mg/kg of selective NOPR antagonist J-113397 or vehicle (i.p.) 30 minutes prior to the session. During the session, mice were presented with 15 trials were a 5s tone preceded 5s of access to a 10% sucrose solution, with a 5-minute null ‘baseline’ period at the beginning and end of the recording. **c**. Averaged traces of NOPLight fluorescence following pre-treatment with vehicle (green) or J-113397 (blue). Traces are aligned to the start of the 5s tone (magenta, shaded) that precedes 5s of access to a 10% sucrose solution (brown, shaded). Data represented as mean ± SEM. **d**. Average number of licks made during sessions that followed vehicle (green) or J-113397 (blue) pre-treatment (two-tailed Wilcoxon test, ns: not significant, n = 7 mice). **e**. Heat map of NOPLight fluorescence corresponding to the average traces in **c** for vehicle (left, green) and J-113397 (right, blue) sessions. **f**. Area under the curve (AUC) for the averaged traces from **c**, calculated over 5-second intervals before/during/after cued-sucrose events. NOPLight fluorescence in the VTA is significantly decreased during and immediately after cued access to 10% sucrose solution (two-tailed Wilcoxon test, **p<0.01, n = 7 mice). Pre-treatment with 20 mg/kg J-113397 blocks this decrease in NOPLight signal (two-tailed Mann-Whitney test, ## p<0.01, n = 7 mice). Data represented as mean ± SEM. **g**. Top: Schematic of fibre photometry setup. Middle: Coronal brain cartoon depicting fibre implant with viral injection of either DIO-GCaMP6m (dark green), FLEX-NOPLight (light green), or NOPLight-ctr (gray) into the VTA of Pnoc-Cre (n = 4), Oprl1-Cre (n = 3), or WT (n = 8) mice, respectively. Bottom: Trial structure for tail lift sessions. **h**. Averaged trace of pnVTA^Pnoc^ GCaMP6m activity during a 10s tail lift. Data represented as mean ± SEM. **i**. Area under the curve (AUC) for photometry trace from **h**, calculated over 5-second intervals before/during/after tail lift events (two-tailed Wilcoxon test, ***p<0.001, ****p<0.0001, n = 4 mice). Data represented as mean ± SEM. **j**. Averaged trace of FLEX NOPLight (green) and NOPLight-ctr (gray) fluorescence during a 10s tail lift. Data represented as mean ± SEM. **k**. Area under the curve (AUC) for photometry traces from **j**, calculated over 5-second intervals before/during/after tail lift events. FLEX-NOPLight fluorescence increases during the tail lift (two-tailed Wilcoxon test, **p<0.01, n = 3 mice). NOPLight-ctr fluorescence does not change during the tail lift (two-tailed Mann-Whitney test, # p<0.05, n = 3-8 mice). Data represented as mean ± SEM.

The wild-type receptor of NOPLight, NOPR, is known to respond highly selectively to N/OFQ, as compared to all other endogenous opioid peptides. ^3,49^ To ensure that NOPLight retained a similar degree of ligand-selectivity, we tested its response to a series of opioid peptides applied to NOPLight-expressing cells at a high concentration (1 μM). NOPLight showed negligible response to dynorphins, enkephalins and β-endorphin (**Fig. 1i**). Similarly, the sensor showed minimal response to a panel of fast neurotransmitters (**Fig. 1j**), indicating high N/OFQ ligand selectivity at NOPLight.

To ensure minimal interference of NOPLight with cellular physiology, we investigated the putative coupling of the sensor with downstream intracellular signalling pathways and compared it to that of wild-type human NOPR. Like other members of the opioid receptor family, NOPR is Gi/o coupled and inhibits basal and Gs-stimulated adenylate cyclase activity upon activation, thus lowering intracellular cAMP levels ^3,49^. We used the GloSensor cAMP assay in HEK293 cells expressing either wild-type human NOPR or NOPLight to monitor intracellular cAMP level with a bioluminescence readout. Application of 1 nM N/OFQ significantly inhibited the forskolin-induced cAMP response in cells expressing the NOPR, while no effect was observed for NOPLight-expressing cells treated with up to 100 nM N/OFQ (**Supplementary Fig. 6a**). At higher concentrations of N/OFQ, inhibition of the cAMP signal in NOPLight-expressing cells was still significantly reduced compared to that of NOPR.

**FIGURE 6.**
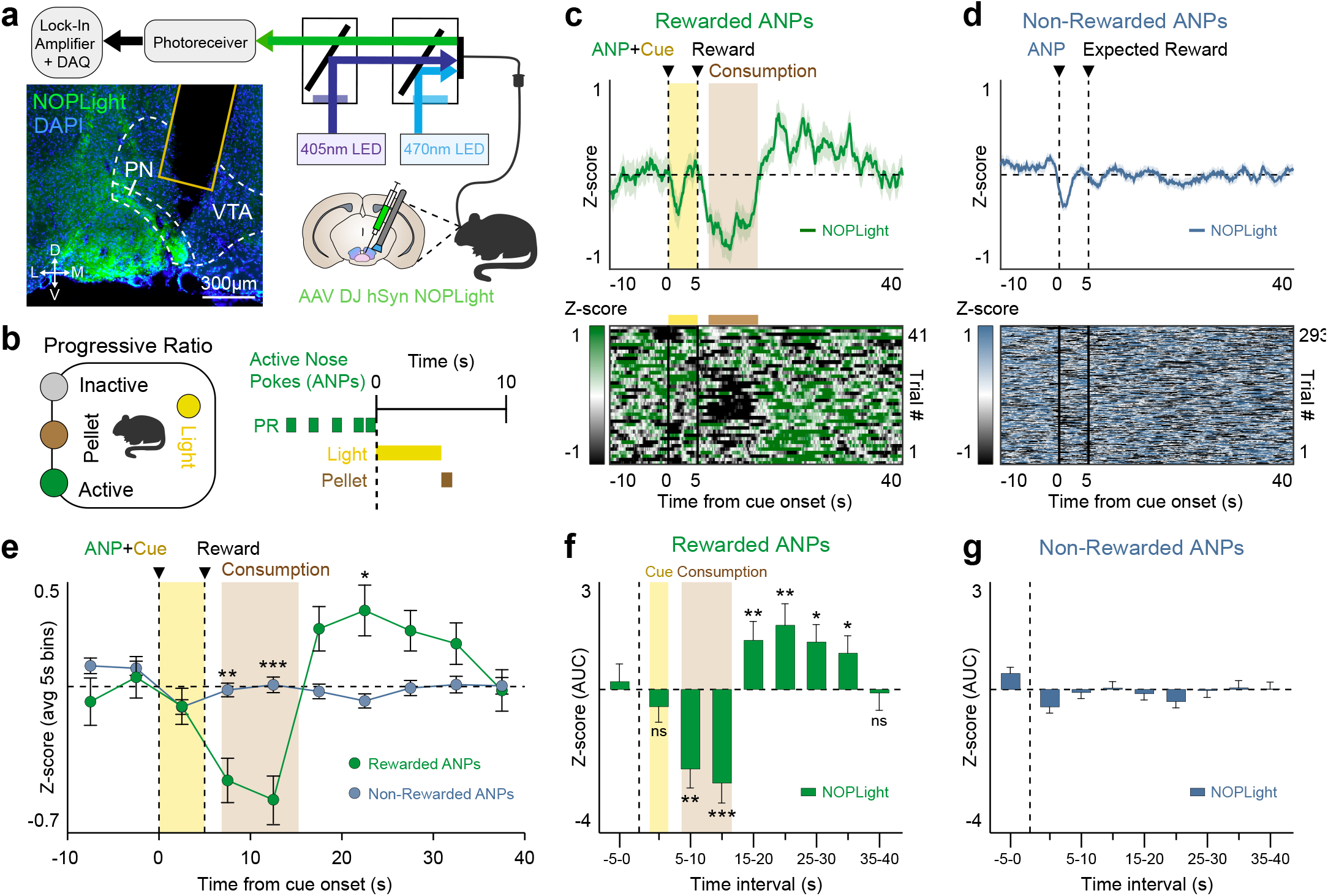
NOPLight detection of endogenous VTA N/OFQ release during high effort reward-seeking. **a**. Schematic of fibre photometry setup. Coronal brain cartoon of viral injection of NOPLight and fibre implant in the VTA of WT mice (n = 4). Representative image showing expression of DAPI (blue) and NOPLight (green) with fibre placement in VTA. **b**. Left: Cartoon depicting operant box setup for the progressive ratio (PR) test. Right: Trial structure for sucrose pellet delivery during the progressive ratio paradigm. **c**. Top: Averaged trace of NOPLight signal for all mice (n = 4), aligned to rewarded active nose pokes (ANPs) made during the PR test. Only active nose pokes that resulted in reward delivery are included. Epoch shown includes the 5s light cue (yellow, shaded) that precedes reward delivery. Time to pellet retrieval and duration of consumption period averaged across all animals (brown, shaded). Data represented as mean ± SEM. Bottom: Corresponding heat map, where each row corresponds to an individual, reinforced ANP epoch. **d**. Top: Averaged trace of NOPLight signal for all mice (n = 4), aligned to non-rewarded ANPs made during the PR test. Data represented as mean ± SEM. Bottom: Corresponding heat map, where each row corresponds to an individual, non-reinforced ANP epoch. **e**. NOPLight fluorescence averaged over 5-second intervals for rewarded (green) and non-rewarded (blue) ANP epochs. Time to pellet retrieval and duration of consumption period averaged across all rewarded trials (brown, shaded). In comparison to the relatively stable NOPLight signal detected across non-rewarded ANP trials, NOPLight signal across rewarded ANP trials decreases during reward consumption then increases immediately post-consumption (two-way repeated-measures ANOVA with Bonferroni’s post hoc test, *p<0.05, **p<0.01, ***p<0.001, n = 4 mice). Data represented as mean ± SEM. **f-g**. Area under the curve (AUC) for photometry traces from **c** and **d** respectively, calculated over 5-second intervals before/during/after rewarded (**f**) and non-rewarded (**g**) active nose pokes (two-tailed Wilcoxon test, *p<0.05, **p<0.01, ***p<0.001, n = 4 mice). Data represented as mean ± SEM.

Under physiological conditions, activation of GPCRs can induce β-arrestin recruitment and/or receptor internalization. We monitored β-arrestin-2 recruitment to the cell surface upon N/OFQ stimulation using TIRF microscopy. Activation of NOPR induced strong β-arrestin-2 recruitment and subsequent internalization of the receptor, whereas NOPLight showed neither of these effects after prolonged occupancy by N/OFQ (**Supplementary Fig.6b-e**). In accordance with the lack of coupling to β-arrestin-2, we also observed that the NOPLight response remained stable for over 1.5 hours in the presence of N/OFQ, and the increase in fluorescence could be reversed by treating the cells with a membrane-impermeable peptide NOPR antagonist (UFP-101, 100 nM; **Supplementary Fig. 6f-g**). These results indicate that while NOPLight retains the ligand selectivity of the native receptor, its cellular expression has a very low likelihood of interfering with intracellular signalling, making the sensor suitable for utilization in physiological settings.

### *Ex vivo* characterization of NOPLight

We then expressed the sensor directly in brain tissue by injecting NOPLight-encoding adeno-associated virus (AAVDJ-hSyn-NOPLight) in the arcuate nucleus (ARC) of the hypothalamus of wild-type mice. After 4 weeks of incubation, the sensor was clearly expressed and was characterized for its functional response to exogenously perfused N/OFQ (**Fig. 2a-b**). NOPLight responses could be detected from ROIs in the ARC upon superfusion of as little as 10 nM N/OFQ on the slice (⍰F/F_0_ = 1.9 ± 1.1%, n = 10), and the magnitude of the responses continued to increase up to the highest N/OFQ concentration tested (10 μM, ⍰F F/F_0_ = 73 ± 21%) (**Fig. 2c**). To measure NOPlight OFF kinetics in brain tissue, we locally puffed N/OFQ next to the sensor-expressing area. The sensor off kinetics ranged between 30-60 s under these conditions (30.9 ± 4.5 s, 41.1 ± 9.9 s and 53.1 ± 6.6 s at 10, 50 and 500 nM respectively, mean ± SEM) (**Supplementary Fig. 7a-d**).

We next tested whether the *in situ* sensitivity of NOPLight would be sufficient to detect endogenous N/OFQ release in this *ex vivo* setting. For this purpose, we again used the preparation of ARC neurons with NOPLight expression and additionally expressed the activating DREADD hM3D in PNOC neurons (**Fig. 2d, e**). For this, a hM3Dq-encoding adeno-associated virus (AAV8-hSyn-DIO-hM3Dq-mcherry) was injected into the ARC of PNOC-Cre mice, enabling Cre-dependent expression of hM3Dq. hM3Dq activation by bath-applied clozapine N-oxide (CNO) evoked a clear increase in the firing rate of PNOC neurons that lasted for several minutes (**Fig. 2f**). Correspondingly, we could detect an increase in NOPLight responses with a mean fluorescence change of ⍰F F/F_0_ = 1.6 ± 0.3% (**Figure 2g-i**), indicating that the sensor could report endogenous N/OFQ release under these conditions. The response of NOPLight was reversible and reflected the time course of chemogenetic activation of PNOC neurons. Overall, these results indicate that NOPLight provides a sensitive and specific readout of both superfused as well as endogenous N/OFQ peptide release in brain tissue.

### NOPLight activation by an exogenous NOPR agonist *in vivo*

Our results *in vitro* (**Fig. 1g**) showed potent and efficacious NOPLight responses to the small molecule NOPR agonist Ro-64. Thus, we determined whether we could use fibre photometry to record NOPLight fluorescence *in vivo* and track the target engagement of this NOPR agonist action in real-time within the brain (**Fig 3**). To do so, we injected WT mice with AAV-DJ-hSyn-NOPLight either in the VTA or in the ARC and implanted optic fibres above the injection sites for photometry recordings (**Fig. 3a-b, Supplementary Fig. 8**). At 3-4 weeks post-viral injection, we detected robust, dose-dependent increases in NOPLight fluorescence in both brain areas following systemic (i.p.) injection of increasing doses of Ro-64 (**Fig. 3c, Supplementary Fig. 8a-d**). To determine if the observed increase in NOPLight fluorescence in the VTA was produced by the sensor binding to the NOPR agonist, we pre-treated animals with two different selective NOPR antagonists, LY2940094 (LY, 10 mg/kg o.g.) or J-113397 (J11, 10 mg/kg i.p.) 30 min prior to injection of Ro-64. Both NOPR-selective antagonists fully inhibited the agonist-induced fluorescent signal (**Fig. 3d-e**).

Next, we injected additional cohorts of OPRL1-Cre (line characterized in the VTA in **Supplementary Fig. 9a-h**) or WT mice in the VTA with AAVs containing either a Cre-dependent variant of NOPLight (AAV-DJ-hSyn-FLEX-NOPLight), or the control sensor (AAV-DJ-hSyn-NOPLight-ctr) respectively, with optic fibres implanted above the injection site (**Fig. 3f**). We found that systemic injection of 10 mg/kg Ro-64 produced a robust increase in FLEX-NOPLight signal that was not significantly different from the agonist-induced signal we recorded from non-conditionally expressed NOPLight (**Fig. 3g-h**). In contrast, the control sensor showed no fluorescent response to a 10 mg/kg Ro-64 injection (**Fig. 3g-h**). These results indicate that NOPLight expressed in freely moving animals reliably provides dose-dependent and antagonist-sensitive detection of exogenous NOPR agonists in real time.

### NOPLight detection of chemogenetically-evoked endogenous N/OFQ release in the VTA

A primary goal motivating our development of the NOPLight sensor was to ultimately achieve real-time detection of local N/OFQ release in behaving animals. Thus, we set out to determine whether NOPLight reliably detects endogenous release of N/OFQ. To accomplish this, we evoked endogenous N/OFQ release in a local paranigral VTA (pnVTA) circuit we previously identified ^22^ by using a chemogenetic approach to selectively activate VTA^PNOC^ neurons while simultaneously recording changes in VTA-NOPLight fluorescence via fibre photometry. PNOC-Cre mice were co-injected in the VTA with two Cre-dependent AAVs containing i) AAV-DH-hSyn-NOPLight and ii) an mScarlet-tagged stimulatory hM3Dq DREADD (AAV5-EF1a-DIO-HA-hM3D(Gq)-mScarlet), with optic fibres implanted above the injection site (**Fig. 4a**). Control animals received an mCherry (AAV5-EF1a-DIO-mCherry) injection in place of the red fluorophore-tagged DREADD. Based on our earlier spectral characterization which showed that the isosbestic point of NOPLight is closer to 440 nm (**Supplementary Fig. 3**), we tested and characterized an alternative set of LED wavelengths in this group of experiments, using 435 nm and 490 nm as the isosbestic and signal wavelengths, respectively. Activation of the hM3Dq DREADD via systemic injection with 5 mg/kg of clozapine-N-oxide (CNO) produced a significant increase in NOPLight fluorescence that was not observed in the control animals expressing mCherry in place of the DREADD (**Fig. 4b**). To confirm that this increase was truly the result of chemogenetically evoked endogenous N/OFQ release and thus acting via a NOPR-dependent mechanism. Pre-treatment with the selective NOPR antagonist LY2940094^50^ (LY, 10 mg/kg, o.g.) 30 min prior to CNO injection prevented the CNO-induced increase in NOPLight fluorescence (**Fig. 4c**). Together, these results provide strong evidence that NOPLight can detect evoked endogenous release of N/OFQ in freely moving animals.

### NOPLight detects dynamics in endogenous N/OFQ VTA tone during head-fixed consummatory behaviors *in vivo*

We also examined NOPLight’s ability to report transient, endogenous N/OFQ release evoked by different naturalistic behavioral states. N/OFQ and its receptor NOPR have been implicated in neuromodulation of a wide variety of essential behavioral processes including stress, aversion, motivation, reward seeking, and feeding. ^22,23,27,51,52,53,54,55,56^ We have previously identified a role for VTA N/OFQ signaling in reward-related and aversive behavior, and using GCaMP found that pnVTA^PNOC^ neurons activity is suppressed during sucrose consumption. ^22^ Therefore, we injected WT mice with AAV-DJ-hSyn-NOPLight in the VTA, implanted optical fibres above the injection site, and secured stainless steel rings to allow for head-fixed fibre photometry recording of NOP-Light during sucrose consumption (**Fig. 5a**). Mice were food-restricted (85-90% of their starting body weight) and then NOPLight signal was recorded during cued access to 10% sucrose solution (15 trials, 5s sucrose access/trial) (**Fig. 5b**). Sucrose trials were cued by a 5-second auditory tone (4 kHz, 80 dB) that preceded the 5 seconds of sucrose access to determine if NOPLight would respond to any salient stimulus rather than specifically to sucrose consumption.

We found that NOPLight fluorescence remained unchanged during the 5-second tone, but significantly decreased during sucrose consumption (**Fig. 5c-f**). In contrast, animals that were pre-treated with the selective NOPR antagonist J-113397 (J11, 20 mg/kg, i.p.) 30 minutes prior to cued sucrose recordings had no change in NOPLight signal during sucrose consumption (**Fig. 5c, f**). Mice made a similar number of licks on the sucrose spout during J-113397 and vehicle sessions, suggesting that the lack of signal change during sucrose consumption following J-113397 treatment was a result of the NOPR antagonist blocking NOP-Light’s detection of endogenous N/OFQ levels. This result is consistent with our previous study, which showed that nociceptin-containing neurons have lower activity during reward consumption. These results demonstrate that NOPLight is sensitive to changes in the endogenous N/OFQ tone *in vivo* within the VTA, in behaviorally relevant appetitive contexts.

### NOPLight detects natural endogenous N/OFQ release following aversive stimuli *in vivo*

In addition to having a well-established role in reward processing, the VTA is known to mediate aversive states. Given that N/OFQ and NOPR are also widely implicated in stress and aversive responses, we evaluated whether NOPLight could detect endogenous N/OFQ release in the VTA during an aversive behavior in freely moving animals. Our previous findings showed that stimulating pnVTA^PNOC^ neurons drives aversive responses, so we predicted that the VTA N/OFQ system would be engaged by an aversive stimulus. To test this, we used fibre photometry to record pnVTA^PNOC^ activity in response to tail suspension (**Fig. 5g**). PNOC-Cre mice injected with AAVDJ-EF1a-DIO-GCaMP6m and implanted with optic fibres in the VTA underwent four trials of a 10-second tail suspension, resulting in a robust increase in GCaMP6m fluorescence lasting for the duration of the suspension (**Fig. 5h-i**). To determine if the increase in pnVTA^PNOC^ calcium activity was reflected by NOPLight as an increase in N/OFQ release in the VTA, we repeated the tail suspension test in OPRL1-Cre mice injected with AAV-DJ-hSyn-FLEX-NOPLight and WT mice injected with AAV-DJ-hSyn-NOPLight-ctr (**Fig. 5g**). Fibre photometry recordings of FLEX-NOPLight showed an increase in fluorescence during tail suspension that was significantly elevated in comparison to NOPLight-ctr (**Fig. 5j-k**). These data indicate that NOP-Light can reliably reports the endogenous release of N/OFQ during aversive behavioral events within the VTA.

### Monitoring endogenous N/OFQ dynamics in reward-seeking operant behavior

Our prior work extensively characterized the calcium activity of pnVTA^PNOC^ neurons during operant responding for reward and identified a role for VTA N/OFQ signaling in constraining motivation to obtain a reward. Therefore, we sought to determine whether our sensor could report the dynamics of endogenous N/OFQ release in the VTA during operant reward-seeking behavior. WT mice injected with AAV-DJ-hSyn-NOPLight and implanted with optic fibres in the VTA were food-restricted (85-90% of their starting body weight) and then trained on a Pavlovian and operant conditioning schedule (**Fig. 6a**). Mice were first trained in Pavlovian conditioning sessions to associate a 5-second house light cue (CS) with delivery of a sucrose pellet (US) (**Supplementary Fig. 10a**). Next, they were trained on a fixed ratio (FR) operant schedule, learning first to perform one (FR1) and later three (FR3) nose pokes into an active port to trigger the light cue and sucrose reward. Mice successfully learned that only nose pokes in the active port would result in reward delivery (**Supplementary Fig. 10b**). Tracking of pellet consumption across the training paradigm also confirmed that mice consumed the majority of rewards they obtained across sessions (**Supplementary Fig. 10c**).

NOPLight fluorescent signals recorded during early Pavlovian conditioning sharply increased in response to onset of the light cue (**Supplementary Fig. 10d**), which is consistent with fibre photometry recordings of the calcium activity (GCaMP6s) of a posterior population of pnVTA^PNOC^ neurons known to project locally within the VTA^22^. Across all Pavlovian and operant conditioning schedules, we observed a robust decrease in NOPLight fluorescence that persisted throughout the reward consumption period, and was immediately followed by a transient increase in signal upon the end of a feeding bout (**Supplementary Fig. 10d-h**). This decrease in signal during reward consumption and subsequent increase at the end of the feeding period is consistent with previously reported calcium activity patterns of posterior pnVTA^PNOC^ neurons recorded during both Pavlovian and operant conditioning. Importantly, during FR3 recordings when mice performed an active nose poke that did not yet meet the threshold for a reward delivery, we observed an increase in NOPLight fluorescence following the nose poke instead of a decrease, indicating that the decrease is related to reward consumption and not the operant action (**Supplementary Fig. 10i**). Taken together, NOPLight signal in the VTA closely resembles known patterns of pnVTA^PNOC^ neuron GCaMP activity during Pavlovian and operant reward-related behaviors.

After completing FR3 operant training, mice were placed in a progressive ratio (PR) test where the required nose poke criterion increases exponentially with each subsequent reward until the animal reaches a motivational breakpoint where they are unwilling to exert any further effort to obtain a reward (**Fig. 6b**). We extracted fibre photometry epochs surrounding active nose pokes that were reinforced with delivery of a sucrose pellet reward, finding that NOPLight fluorescence decreases during reward consumption in the PR test and transiently increases immediately upon completion of a consumption bout (**Fig. 6c, f**). In contrast, NOPLight signal remained stable during epochs were animals performed an unsuccessful active nose poke that did not result in reward delivery (**Fig. 6d**, **g**). NOPLight signal during reward consumption is significantly suppressed in comparison to time periods where animals expected but did not obtain a reward (**Fig. 6e**). Notably, our detection of negative changes to NOPLight fluorescence relative to baseline suggests the presence of N/OFQ tone in the VTA, which is consistent with our previous findings regarding this circuit. Collectively, these results indicate that NOPLight faithfully reports bidirectional changes in N/OFQ release in the VTA during freely moving behaviors. Our findings demonstrate NOPLight’s utility in detecting endogenous N/OFQ release during discrete behavioral epochs within the VTA and potentially other N/OFQ brain regions.

## DISCUSSION

Here we describe the engineering, characterization, and application of a novel genetically encoded sensor (NOPLight) for monitoring the opioid neuropeptide N/OFQ *in vitro, ex vivo*, and *in vivo*. Endogenous opioid peptides represent one of the largest classes of neuropeptide families, yet detecting their release, dynamics, and properties *in vitro* and *in vivo* has been a challenge for over 60 years since their discovery. ^57,58,59^ We sought to develop a sensor which could detect: 1) evoked release, 2) endogenous release during naturalistic behavior, and 3) exogenous ligands *in vivo* to inform brain localization of pharmacological agents. These properties have been long sought after to better understand neuropeptide transmission generally, and more specifically opioid peptides and their actions.

The neuropeptide biosensor we developed exhibits a large dynamic range both in HEK293T cells and in neurons, sub-second activation kinetics, very high ligand selectivity, a similar pharmacological profile to that of NOPR, and no detectable interference with endogenous signaling pathways. We demonstrated that NOPLight dose-dependently responds to systemic administration of a NOPR agonist, with sensitivity to blockade by selective NOPR antagonists. NOPLight is also capable of detecting endogenous release of N/OFQ evoked by either chemogenetic stimulation (hM3Dq DREADD) of PNOC neurons or during natural behavior.

N/OFQ and NOPR are widely implicated in diverse behavioral states, which was reflected by our fibre photometry recordings of NOP-Light during varied behaviors. Our recordings of NOPLight signal during head-fixed behavior revealed a NOPR antagonist sensitive decrease in N/OFQ signaling time-locked with access to a 10% sucrose solution. We also identified an increase in N/OFQ release in response to an aversive tail suspension using NOPLight, which was not present in recordings of our control variant of the sensor. While it has been suggested that N/OFQ is released in response to aversive or stressful stimuli, our findings are the first to directly detect endogenous N/OFQ release during an aversive response.

Previous GCaMP recordings of PNOC neurons with local input in the VTA during operant conditioning tasks revealed dynamic engagement of pnVTA^PNOC^ neurons during reward-seeking and consumption behavior. ^22^ While in many cases calcium mobilization is required for dense core vesicle fusion, ^60^ calcium activity is not a direct correlate for peptide release and as such the dynamics of released N/OFQ could not be established in this prior study. Here we report NOPLight activity in the VTA in reward-seeking behavior during fixed ratio-1, -3, and progressive ratio paradigms, which identified a rapid and sustained decrease during reward consumption, and a transient increase after consumption had ended (**Fig. 6, Supplementary Fig. 10**). This pattern of NOPLight signal closely resembles the expected dynamics of N/OFQ release based on the prior study’s GCaMP recordings. ^22^ Notably, we observed both dynamic increases and decreases in NOPLight fluorescence during behavioral epochs, suggesting that the NOPLight can be used to detect changes in peptide tone over behaviorally relevant timescales. These data also provide the ability to align neuronal activity measured either by calcium dynamics or electrophysiology with neuropeptide release during freely-moving behavioral epochs. Given the recent discoveries that PNOC and NOPR are important for motivation, feeding, and sleep induction, ^22,23,51,52,53,54^ understanding the dynamic properties of this opioid peptide system is now of even greater importance as this receptor is now considered a major target for insomnia, addiction, and depression. ^50,61,62^

The endogenous activity of PNOC neurons in the VTA is thought to provide inhibitory tone onto VTA dopamine neurons, constraining motivation to seek rewards. ^15,22,24,63,64,65^ Consistent with this concept, here we observed a decrease in NOPLight fluorescence when animals engaged in reward consumption. Additionally, we measured an increase in NOPLight signal immediately upon completion of a reward consumption bout, suggesting that N/OFQ is released in the VTA after animals have consumed an obtained reward. Since NOPR is largely expressed on dopamine neurons in the VTA and exerts inhibitory influence over their activity, ^24,63^ it is possible that this increase in endogenous N/OFQ release after consumption reflects a temporarily satiated state where N/OFQ signaling transiently increases to suppress tonic dopamine neuron activity, thus reducing motivation to seek out additional rewards. Importantly, this is the first detection of real-time N/OFQ release in this context. As a result, these findings provide insight into the dynamics of N/OFQ signaling in the VTA which acts to coordinate motivated behavior through dopaminergic interactions. Future studies will be able to employ NOPLight in concert with the recently developed red-shifted dopamine sensors ^66^ and other N/OFQ-selective tools like the new OPRL1-Cre line (**Supplementary Fig. 9**) to improve our understanding of N/OFQ modulation of dopamine circuitry during reward-seeking.

We tested the performance of the sensor *in vivo* using different wavelength pairs. Following the development of GCaMPs, sensor excitation has conventionally employed a 405 nm wavelength light as a read-out for non-ligand sensitive signals (known as the isosbestic channel), that typically acts as an internal control for fibre photometry experiments, particularly in freely moving animals. ^67,68^ As a result, most commercially-available photometry setups are tailored to accommodate this wavelength. In this work we noted that, based on the results from our spectral characterization of NOPLight, excitation at a wavelength of 435 nm is better suited as a control ‘isosbestic’ channel. It is particularly worth noting that many recently-developed intensiometric GPCR-based biosensors exhibit a similar spectral property with a right-shifted isosbestic point (i.e. > 420 nm). ^46,69,70,71,72^ In these cases, use of 405 nm as the isosbestic channel for these sensors may lead to confounding results or difficulty in interpretation in some contexts, such as regions where N/OFQ release in response to a given behavior is relatively low, or during behaviors that are highly subject to motion artefact. Since it can be cost and time prohibitive for research groups to add an alternative recording parameter to existing photometry setups, we demonstrated that we were still able to detect meaningful changes in NOPLight fluorescence when using the 405 nm wavelength in some of our fibre photometry recordings. Furthermore, head-fixation and red fluorophore-based motion controls are commonly used as alternatives to isosbestic controls that could easily be implemented with NOP-Light. ^68,73^ Our careful evaluation of NOPLight’s performance when recorded with different isosbestic and excitation wavelengths provides valuable insight into its photophysical properties that will help inform successful application of NOPLight and other neuropeptide sensors in future studies. Our work here lays critical groundwork that will need to be built upon through continued use of NOPLight and its control variant in similar reward-related behaviors, different brain regions, and with additional controls, which will be important for optimizing their implementation and developing general best practices amidst the rapidly expanding use of fluorescent peptide sensors.

Our findings present NOPLight as a unique approach to improve investigations of endogenous opioid peptide dynamics with high spatiotemporal resolution. We characterized NOPLight expression, selectivity, and sensitivity to endogenous N/OFQ release both *in vivo* and *in vitro*. Future optimization of the sensor should seek to improve quantum yield (fluorescent readout) at lower peptide concentrations and to develop red-shifted variants to provide more flexibility in multiplexing NOPLight with other optical tools and sensors. This sensor directly helps to address a longstanding limitation in understanding the real-time dynamics of endogenous peptide release during behavioral epochs. Future applications of neuropeptide sensors such as NOPLight will advance our understanding of the underlying mechanisms by which endogenous opioid peptides control, stabilize and modulate neural circuits to regulate behavior.

## Abbreviations

Pnoc: prepronociceptin
N/OFQ: nociceptin/orphanin FQ
VTA: ventral tegmental area
CeA: central amygdala

## AUTHOR CONTRIBUTIONS

TP and MRB led the study. XZ, CS, TP, and MRB conceptualized and designed the study. XZ developed the NOPLight and NOPLight-ctr sensors, performed *in vitro* sensor screening and characterization, including confocal imaging in HEK293T cells and neurons, and performed kinetic measurements in acute brain slices, under the supervision of TP. MAB prepared primary neuronal cultured under supervision of DB. JD performed two-photon spectral characterization under the supervision of LR, TP and BW. KA performed signaling assays under the supervision of MS. DF and PK performed and analysed acute brain slice electrophysiology and imaging experiments. POP, CAB and LS performed and analysed *in vivo* photometry data in arcuate nucleus under supervision of JCB. CS performed and analysed *in vivo* photometry and chemogenetic experiments in the pnVTA under the supervision of MRB. ALP, AS, RDP, ASA, JCJ, SJ helped with surgeries and behavioral animal training for experiments in pnVTA, under the supervision of MRB. TP, MRB, XZ and CS wrote the manuscript with contributions from all authors.

ACKNOWLEDGMENTS

Council (ERC) under the European Union’s Horizon 2020 research and innovation program (Grant agreement No.s 891959 and 101016787 to TP; 742106 to JCB). We also acknowledge funding from the University of Zürich Forschungskredit (FK-20-042, XZ), the Swiss National Science Foundation (Grant No. 310030_196455, TP; PCEFP3_181282, MS), the NIMH P50MH119467 (MRB), NIH/NIMH F31 F31DA059438-01A1 (CAS), and the Deutsche Forschungsgemeinschaft (401832153, 431549029, EXC 2030-390661388, P.K). We would like to thank Jean-Charles Paterna and the Viral Vector Facility of the Neuroscience Center Zürich (ZNZ) for the kind help with virus production. Some figure panels were created using BioRender.com.

## DATA AVAILABILITY

DNA and protein sequences for the new sensors developed in this study have been deposited on the National Center for Biotechnology Information database (accession nos. OQ067483 and OQ067484) and are available in Supplementary Data S1. DNA plasmids used for viral production have been deposited both on the UZH Viral Vector Facility (https://vvf.ethz.ch/) and on Addgene. Viral vectors can be obtained either from the Patriarchi laboratory, the UZH Viral Vector Facility, or Addgene. All source data are provided with the manuscript. All other raw data can be made available upon reasonable request.

## CODE AVAILABILITY

Custom MATLAB code is available on GitHub at https://github.com/patriarchilab/NOPLight.

## SUPPORTING INFORMATION

Additional supporting information may be found in the online version of the article.

## MATERIALS AND METHODS

### Molecular cloning and structural modelling

The prototype sensor was designed *in silico* by sequence alignment (Clustal Omega2) and ordered as a geneblock (Thermo Fisher) flanked by HindIII and NotI restriction sites to be subsequently cloned into pCMV vector (Addgene #111053). For sensor optimization, site directed mutagenesis and Circular Polymerase Extension Cloning was performed by polymerase chain reaction with custom designed primers using a Pfu-Ultra II fusion High Fidelity DNA Polymerase (Agilent). Sanger sequencing (Microsynth) was performed for all constructs reported in the manuscript. The structural prediction of the NOPLight was generated by a deep learning-based modelling method, RoseTTAFold. ^4^

### Cell culture, confocal imaging and quantification

HEK293T cells (ATCC CRL-3216) were authenticated by the vendor. They were seeded in glass bottom 35-mm (MatTek, P35G-1.4-14-C) or 24-well plates (Cellvis, P24-0-N) and cultured in Dulbecco’s modified Eagle’s medium (Gibco) with 10% Fetal Bovine Serum (Gibco) and antibiotic-antimycotic (1:100 from 10,000 units/ml penicillin; 10,000 μg/ml streptomycin, 25 μg/ml amphotericin B, Gibco) mix at 37°C and 5% CO2. Cells were transduced at 70% confluency using Effectene transfection kit (QIAGEN) and imaged after 24 – 48 hours. Primary neuronal cultures were prepared as the following: the cerebral cortex of 18 days old rat embryos was carefully dissected and washed with 5 ml sterile-filtered PBGA buffer (PBS containing 10 mM glucose, 1 mg/ml bovine serum albumin and antibiotic-antimycotic 1:100 (10,000 units/ml penicillin; 10,000 μg/ml streptomycin; 25 μg/ml amphotericin B)). The cortices were cut into small pieces with a sterile scalpel and digested in 5 ml sterile filtered papain solution for 15 min at 37°C. The supernatant was removed, and tissue washed twice with complete DMEM/FCS medium (Dulbecco’s Modified Eagle’s Medium containing 10% Fetal Calf Serum and penicillin/streptomycin, 1:100). Fresh DMEM/FCS was then added, and the tissue gently triturated and subsequently filtered through a 40-μm cell-strainer. Finally, the neurons were plated at a concentration of 40,000-50,000 cells per well onto the poly-L-lysine (50 μg/ml in PBS) coated 24-well culture plate and incubated overnight at 37°C and 5% CO_2_. After 24 hours of incubation, the DMEM medium was replaced with freshly prepared NU-medium (Minimum Essential Medium (MEM) with 15% NU serum, 2% B27 supplement, 15 mM HEPES, 0.45% glucose, 1 mM sodium pyruvate, 2 mM GlutaMAX). Cultured neurons were transduced at 4 days *in vitro* (DIV4) with AAV-DJ-hSynapsin1-NOPLight, AAV1-hSyn1-NES-jRCaMP1b, or AAV-DJ-hSynapsin1-NOPLight-ctr viruses in culture media using a final titer for each virus between 4x10^9^ and 4x10^10^ GC/ml, and were imaged between DIV19–21. All reagents used are from Gibco. Unless otherwise noted, confocal imaging for all constructs reported in the manuscript are performed as follows:

Images were acquired on an inverted Zeiss LSM 800 microscope with a 488-nm laser for NOPLight and NOPLight-ctr, and a 564-nm laser for Red-Dextran dye and JRCaMP1b. For characterization of the dynamic range, expression level and pharmacological properties, HEK293T cells and/or neurons expressing the construct were first rinsed with HBSS (Gibco) and imaged at a final volume of 100 μL HBSS under a 40x objective. For testing sensor performance *in vitro* in various pH, NOPLight expressing HEK293T cells were first incubated in PBS (Gibco) with adjusted pH (6 - 8) for 3 minutes before addition of the ligand. For pharmacological characterizations, the following compounds were used: Nocistatin (Abbiotec); J-113397 (Sigma-Aldrich); UFP-101 (Sigma-Aldrich); Ro 64-6198 (Sigma-Aldrich); MCOPPB (Cayman); Orphanin FQ (1-11) (Tocris); Leu-Enkephalin (Cayman); Met-Enkephalin (Cayman); Dynorphin A (Cayman); Dynorphin B (Cayman); β-Endorphin (Sigma-Aldrich); γ-Aminobutyric acid (Sigma-Aldrich); Dopamine hydrochloride (Sigma-Aldrich), Acetylcholine bromide (Sigma-Aldrich), Glutamate (Sigma-Aldrich). All compounds are diluted to the desired final concentration in HBSS before experiment except Ro 64-6198 and J-113397, which were diluted in <0.02% DMSO. All ligands were carefully pipetted into the imaging buffer during experiment. To determine the apparent affinity of the sensor, HEK293T cells and neurons cultured in glass bottom 24-well plates were rinsed with HBSS and imaged under a 20x objective with a final buffer volume of 500 μL HBSS. Ligands were manually applied on the cells during imaging to reach the desired final concentration.

ΔF/F_0_ was determined as the ratio of change in fluorescence signal change upon ligand activation and the baseline fluorescence level

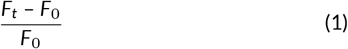

where F_0_ is determined as the mean intensity value over the baseline imaging period.

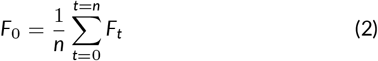

Unless stated otherwise, only pixels corresponding to cell membrane were considered as regions of interest (ROIs) thus included in the analysis. ROIs were selected by auto-thresholding function of ImageJ and confirmed by visual inspection.

### Kinetic measurements and analysis

To obtain the time constant for sensor activation, red fluorescent dye Antonia Red-Dextran (3000 M.W., Sigma-Aldrich) and N/OFQ (MedChem Express) were simultaneously applied in bolus to sensor-expressing HEK293T cells at 37°C with a stage-top incubator (Tokai Hit). Fluorescent signals were excited at 488 nm (NOPLight) and 561 nm (Red-Dextran dye) and recorded using the high-speed line-scan function (Zeiss LSM 800) at 800Hz. The onset latency of each experiment was first determined by calculating the time for the red-dextran fluorescent signal to reach 85% percent of maximal value at the plateau. Only experiments with an onset latency smaller than 50 ms were considered in subsequent analysis to minimize the contribution of N/OFQ peptide diffusion to the temporal profile of sensor response. Membrane-corresponding pixels were first selected by thresholding pixel-wise ΔF/F_0_ at 65% criteria. Fluorescent signal change of each membrane pixel was then normalized and fitted by a mono-exponential association model using custom-written MATLAB script to derive the time constant τ^on^.

To obtain the time constant for ligand wash-off, HEK293T cells were cultured on Poly-D-Lysine (Gibco) coated 18-mm coverslips in Dulbecco’s modified Eagle’s medium (Gibco) with 10% Fetal Bovine Serum (Gibco) and antibiotic-antimycotic (1:100 from 10,000 units/ml penicillin; 10,000 μg/ml streptomycin, 25 μg/ml amphotericin B, Gibco) mix at 37°C and 5% CO2. Cells were transduced at 70% confluency using Effectene transfection kit (QIAGEN) and imaged after 24 – 48 hours. On the day of recording, coverslips with NOPLight expressing HEK293T cells were transferred to the imaging chamber perfused with 37°C HBSS (Gibco). Fluorescence response was obtained with a blue (469 nm) LED (Colibri 7, Zeiss) on an upright Axio Examiner A1 microscope (Zeiss) using an N-Achroplan 10x/0.3 M27 objective (Zeiss). Images were collected at a sampling rate of 1 Hz (Live Acquisition, Thermo Fisher Scientific). For wash-off experiments, N/OFQ was added into the imaging chamber to a final concentration of 500 nM with the perfusion flow turned off. The perfusion system was turned on after 30 s of ligand application to allow equilibrium of sensor response. For puffed application experiments, a glass pipette mounted on a microinjector (Nanoject II™ Drummond) filled with N/OFQ was place near the imaged field of view. 50 nL of 100 μM N/OFQ was puffed onto the cells at a rate of 50 nL/s. Regions of interest for quantification of the fluorescence response was selected in Fiji (Image J). All cell membranes in the field of view were selected for analysis for wash-off experiments, whereas in puffed application experiments, only cell membranes within the radius of ½ of the glass pipette diameter were selected. A mono-exponential decay model was fitted to the data using Curve Fitting Toolbox (Matlab) to deduce the time constant τ _off_.

### One-photon spectral characterisation

One-photon fluorescence excitation (l^em^ = 560 nm) and emission (l^exc^ = 470 nm) spectra were determined using a Tecan M200 Pro plate reader at 37°C. HEK293T cells were transfected with Effectene transfection kit (QIAGEN). 24 hours after transfection cells were disscociated with Versene (Thermo Fisher) and thoroughly washed with PBS. Next, cells were resuspended in PBS to a final concentration of 3.3 x 10^6^ cells/mL and aliquoted into two individual wells of a 96-well microplate with or without ligand (N/OFQ or Ro 64-6198, 1 μM), together with two wells containing the same amount of non-transfected cells to account for autofluorescence and a single well containing PBS to determine the Raman bands of the solvent. To determine the excitation and emission spectra of NOPLight at various pH, PBS was adjusted to the desired pH using 10 M HCl or 10 M NaOH.

### Two-photon brightness characterisation

Two-photon brightness profiles of NOPLight were obtained from HEK293T cells before and after addition of N/OFQ (1 μM). Cells were transfected with Lipofectamine 3000 and were imaged 24 hours post transfection. The medium was replaced with PBS prior to imaging in order to avoid DMEM autofluorescence. The two-photon spectra were acquired as described previously. ^46^

### cAMP assay

HEK293 cells growing at 70% confluency in a 10-cm dish were transfected with wild-type human NOPR or NOPLight (3 μg DNA) and GloSensor-20F (Promega, 2.5 μg DNA) using 12 μL Lipofectamine 2000 (Thermo Fisher) as in. ^74^ After 24 h, cells were plated into clear-bottom 96-well plates at 200,000 cells/well in DMEM (without phenol red, with 30 mM HEPES, p.H. 7.4) containing 250 μg/ml luciferin and incubated for 45-60 min at 37°C. Cells were treated with forskolin (3 μM) and varying concentrations of N/OFQ immediately followed by image acquisition every 45 s for 30 min at 37°C using a Hidex Sense plate reader. Luminescence values were normalized to the maximum luminescence values measured in the presence of 3 μM forskolin and to vehicle treated control cells.

### TIRF microscopy

HEK293 cells growing on polylysine-coated 35-mm, glass-bottom dishes (MatTek, P35G-1.5-14-C) were transfected with wild-type human NOPR or NOPLight (0.8 μg DNA) and β-arrestin-2-mCherry (1 μg DNA) using 3 μL Lipofectamine 2000 (Thermo Fisher). After 24 h, receptors were surface-labeled for 10 min with anti-FLAG M1-AF647^46^ and media changed to HBS imaging solution (Hepes buffered saline (HBS) with 135 mM NaCl, 5 mM KCl, 0.4 mM MgCl_2_,1.8 mM CaCl_2_, 20 mM Hepes, 5 mM d-glucose adjusted to pH 7.4). Cells were imaged at 37°C using a Nikon TIRF microscope equipped with a 100x 1.49 oil CFI Apochromat TIRF objective, temperature chamber, objective heater, perfect focus system and an Andor DU897 EMCCD camera, in time-lapse mode with 10 s intervals. The laser lines used were 561 nm (for β-arrestin-2) and 647 nm (for receptor constructs). 10 μM N/OFQ was added by bath application. Protein relocalization (F) was calculated as F(t)/F0 with F(t) being the β-arrestin-2 signal at each time point (t) (normalized to M1-AF647 signal, when specified) and F0 being the mean signal before ligand addition.

### Virus production

The adeno associated virus (AAV) encoding NOPlight was produced by the Viral Vector Facility at the University of Zurich (VVF). The AAVs encoding the NOPlight-ctr sensor and the Cre-dependent NOPlight were produced by Vigene Biosciences. All other viruses used in this study were obtained either from the VVF or Addgene. The viruses used in this study were: AAVDJ-hSyn-NOPLight, 4.1x10^13^ GC/ml; AAVDJ-hSyn-NOPlight-ctr, 2.9x10^13^ GC/ml; AAVDJ-hSyn-FLEX-NOPlight, 2.5x10^13^ GC/ml; AAV5-EF1a-DIO-HA-hM3D(Gq)-mScarlet, 1.7x10^13^ GC/ml; AAV5-DIO-hSyn-mCherry, 2.4x10^12^ GC/ml; AAV8-hSyn-DIO-hM3Dq.mcherry, 1x10^13^ GC/ml; AAV-DJ-EF1a_DIO-GCaMP6m 1.2x10^13^ GC/ml; AAV1-hSyn1-NES-jRCaMP1b, 6.9x10^12^ GC/ml.

### Animals

Ten-to 24-week-old, wild-type C57BL/6 mice, PNOC-Cre (as described previously ^22,23^ and OPRL1-Cre mice were used in this study. Animal procedures were performed in accordance to the guidelines of the European Community Council Directive or the Animal Welfare Ordinance (TSchV 455.1) of the Swiss Federal Food Safety and Veterinary Office and were approved by the Zürich Cantonal Veterinary Office, and the other respective local government authorities (Bezirksregierung Köln, Animal Care and Use Committee of University of Washington). Mice were housed at the animal facility at 22–24°C and a 12 h/12 h light/-dark cycle and ventilated cages receiving a standard chow and water ad libitum. Animals placed in the Pavlovian and operant conditioning paradigms were food restricted down to 90% of their ad libitum body weight beginning one-week prior to conditioning for the entire duration of the paradigm.

### Stereotaxic surgery

#### Viral vector injections and optic fibre implantation in the arcuate nucleus (ARC)

All surgeries were performed on male adult mice aged 8-10 weeks. Animals were anesthetized with 5% isoflurane and maintained at 1.5-2% throughout the surgery. For cell-specific DREADD expression, adeno-associated virus encoding AAV8-hSyn-DIO-hM3Dq.mcherry (Addgene #44361, 100 nL, viral titer 1 x 10^13^ GC/mL) was injected using a glass capillary into the ARC (−1.5 mm AP, -0.3 mm ML, -5.78 mm DV). Adeno-associated virus encoding AAV-DJ-hSyn-NOPLight (500 nL, viral titer 0.8 x 10^13^ GC/mL) was injected into the ARC in either wild-type or DREADD-expressing PNOC-Cre mice. All viruses were injected at a rate of 100 nL/min. For *in vivo* fibre photometry, after 5 min of viral injections a sterile optic fibre was implanted (diameter 400 μm, Doric Lenses) at the following coordinates (−0.450 mm AP, -0.2 mm ML, -5.545 mm DV) at an 8° angle. The implant was fixed with dental acrylic and closed with a cap after drying. Mice were allowed to recover from surgery 4 weeks before *in vivo* fibre photometry recordings started.

#### Viral vector injections and optic fibre implantation in the VTA

Following a minimum of seven days of acclimation to the holding facility, mice were initially anesthetized in an induction chamber (1-4% isoflurane) and placed into a stereotaxic frame (Kopf Instruments, Model 1900) where anesthesia was maintained at 1-2% isoflurane. Depending on the specific experimental paradigm, mice received viral injections either unilaterally or bilaterally using a blunt neural syringe (86200, Hamilton Company) at a rate of 100nL/min. For exogenous pharmacology experiments, wild-type (WT) mice were injected with AAV-DJ-hSyn-NOPLight (200-500 nL, viral titer 2-4 x 10^12^ vg/mL) or AAV-DJ-hSyn-NOPLight-ctr (300 nL, viral titer 2.9 x 10^13^ vg/mL) and OPRL1*-*Cre mice were injected with AAV-DJ-hSyn-FLEX-NOPLight (300 nL, viral titer 2.5 x 10^13^ vg/mL). For chemogenetic experiments, PNOC*-*Cre mice were co-injected with a 1:1 mix of AAV-DJ-hSyn-FLEX-NOPLight (300 nL, viral titer 2.5 x 10^12^ vg/mL) and either AAV5-EF1a-DIO-HA-hM3D(Gq)-mScarlet (300 nL, viral titer 1.7 x 10^13^ vg/mL) or AAV5-DIO-hSyn-mCherry (300 nL, viral titer 2.4 x 10^12^ vg/mL). For head-fixed behavioral experiments, WT mice were injected with AAV-DJ-hSyn-NOPLight (300 nL, viral titer 4.1 x 10^12^ vg/mL). For freely moving tail lift experiments, PNOC-Cre mice were injected with AAV-DJ-EF1a-DIO-GCaMP6m (300nL, viral titer 1.2 x 10^13^ vg/mL); OPRL1-Cre mice were injected with AAV-DJ-hSyn-FLEX-NOPLight (300nL, viral titer 2.5 x 10^12^ vg/mL); WT mice were injected with AAV-DJ-hSyn-NOPLight-ctr (300nL, viral titer 2.9 x 10^12^ vg/mL). For freely moving reward seeking experiments (Pavlovian conditioning, operant conditioning, progressive ratio test), WT mice were injected with AAV-DJ-hSyn-NOPLight (300 nL, viral titer 4.1 x 10^13^ vg/mL). All viruses were injected into the VTA (stereotaxic coordinates from Bregma: -3.3 to -3.4 AP, +1.6 ML, -4.75 to -4.3 DV) at a 15° angle, with an optic fibre implanted above the injection site. For animals undergoing head-fixed behaviors, a stainless-steel head-ring was also secured to allow for head fixation. All implants were secured using Metabond (C & B Metabond). Animals were allowed to recover from surgery for a minimum of 3 weeks before any behavioral testing, permitting optimal viral expression.

#### Viral vector injections in Nucleus Accumbens

Surgeries were performed on wildtype mice (both male and female) aged 8–10 weeks. Anesthesia was induced using 4–5% isoflurane and maintained at 2%. AAV-DJ-hSyn-NOPLight (300 nL, viral titer 4 x 10^12^ vg/mL) was injected bilaterally (Nanoject II™ Drummond) at +1.5 AP, ±0.7 ML, -4.5 DV. Animals were allowed to recover from surgery for a minimum of 4 weeks before experiment, permitting optimal viral expression.

### Acute brain slice preparation, electrophysiology and imaging

For recordings in the ARC, experiments were performed at least 4 weeks after viral injections. The animals were lightly anesthetized with isoflurane (B506; AbbVie Deutschland GmbH and Co KG, Ludwigshafen, Germany) and decapitated. Coronal slices (280 μm) containing NOP-Light expression in the arcuate nucleus (ARC) were cut with a vibration microtome (VT1200 S; Leica, Germany) under cold (4°C), carbogenated (95% O_2_ and 5% CO_2_), glycerol-based modified artificial cerebrospinal fluid (GaCSF: 244 mM Glycerol, 2.5 mM KCl, 2 mM MgCl2, 2 mM CaCl2, 1.2 mM NaH2PO4, 10 mM HEPES, 21 mM NaHCO3, and 5 mM Glucose adjusted to pH 7.2 with NaOH). The brain slices were continuously perfused with carbogenated artificial cerebrospinal fluid (aCSF: 125 mM NaCl, 2.5 mM KCl, 2 mM MgCl_2_, 2 mM CaCl_2_, 1.2 mM NaH_2_PO_4_, 21 mM NaHCO_3_, 10 mM HEPES, and 5 mM Glucose adjusted to pH 7.2 with NaOH) at a flow rate of 2.5 ml/min. To reduce GABAergic and glutamatergic synaptic input, 10^−4^ M PTX (picrotoxin, P1675; Sigma-Aldrich), 5 x 10^−6^ M CGP (CGP-54626 hydrochloride, BN0597, Biotrend), 5 x 10^−5^ M DL-AP5 (DL-2-amino-5-phosphonopentanoic acid, BN0086, Biotrend), and 10^−5^ M CNQX (6-cyano-7-nitroquinoxaline-2,3-dione, C127; Sigma-Aldrich) were added to perfusion aCSF. The imaging setup consisted of a Zeiss AxioCam/MRm CCD camera with a 1388x1040 chip and a Polychromator V (Till Photonics, Gräfelfing, Germany) coupled via an optical fibre into the Zeiss Axio Examiner upright microscope. NOPLight fluorescence was collected at 470 nm excitation with 200 ms exposure time and a frame rate of 0.1 Hz. The emitted fluorescence was detected through a 500-550 nm bandpass filter (BP525/50), and data were acquired using 5x5 on-chip binning. Images were recorded in arbitrary units (AU) and analyzed as 16-bit grayscale images. To investigate the responses of NOPLight to N/OFQ (cat# 0910, Tocris), increasing concentrations (5 nM, 10 nM, 50 nM, 100 nM, 500 nM, 1 μM, 5 μM, 10 μM) of N/OFQ were sequentially bath-applied for 10 min each. Changes in fluorescent intensity upon ligand application were quantified by comparing the averaged fluorescent intensity measured in 3 min intervals immediately before and at the end of ligand application. For electrophysiological recordings, the preparation of the brain slices and the recording conditions were the same as for the imaging experiments. Perforated patch clamp recordings were performed as previously described. ^23^ To investigate NOPLight response to chemogenetically-evoked endogenous N/OFQ release, hM3Dq was activated by bath application of 3 μM clozapine N-oxide (CNO, ab141704, abcam) for 10 min. N/OFQ and CNO were bath-applied at a flow rate of 2.5 ml/min. The analysis was performed offline using ImageJ (version 2.3.0/1.53f) and Prism 9 (GraphPad, California, USA). Amplitudes and kinetics of the signals were calculated as means (in AU) of fluorescent regions in the ARC, which were defined as the respective regions of interest (ROI, 0.15 – 0.2 mm^2^). Biexponential fits of the signals’ time courses before N/OFQ application were used to correct for bleaching.

For recordings in the NAc, experiments were performed at least 4 weeks after bilateral viral injections. Mice were anesthetized with an intraperitoneal injection of pentobarbital (200 mg/kg, 10 mL/kg) and decapitated. The brain was quickly extracted while submerged in ice-cold aCSF (120 mM NaCl, 2.5 mM KCl, 1.25 mM NaH2PO4, 26 mM NaHCO3, 5 mM HEPES, 1 mM MgCl2, 14.6 mM D-glucose and 2.5 mM CaCl2 at 305-310 mOsm/kg) bubbled with 95/5% O2/CO2. 275-μm thick coronal slices containing the NAc were obtained using a vibratome (HM 650V, ThermoFisher Scientific). The slices were incubated at 34°C for 20 minutes in continuously oxygenated aCSF. Following incubation, brain slices were transferred at RT and kept until recording. Recordings were conducted in a slice chamber kept at 31°C perfused with aCSF. To visualize NOPLight signal, slices were illuminated with a blue (469 nm) LED (Colibri 7, Zeiss) on an upright Axio Examiner A1 microscope (Zeiss) using an N-Achroplan 10x/0.3 M27 objective (Zeiss). Images were collected at a sampling rate of 1 Hz (Live Acquisition, Thermo Fisher Scientific). To locally puff N/OFQ, a glass pipette filled with the desired concentration of N/OFQ was mounted to a microinjector (Nanoject II™ Drummond) and positioned into the imaged field of view on the slice. Various concentrations of N/OFQ (in 50 nL) were puffed onto the slice at 50 nL/s rate. Fluorescence responses were quantified in Fiji (Image J) by selecting a circular region of interest with a radius of ½ of the size of the glass pipette diameter.

### Tissue preparation and immunohistochemistry (IHC)

#### IHC in the ARC

Brain slices were fixed in Roti-Histofix (PO873, Carl Roth) for 12 h at 4°C and subsequently rinsed in 0.1 M phosphate-buffered saline (PBS, 3 x 10 min). PBS contained (in mM) 72 Na_2_HPO_4_ x dihydrate, 28 NaH_2_PO_4_ monohydrate, resulting in pH 7.2. To facilitate antibody penetration and prevent unspecific antibody binding, brain slices were preincubated in PBS containing 1% (w/v) Triton X-100 (TX, A1388, AppliChem) and 10% (v/v) normal goat serum (NGS, ENG9010-10, Biozol Diagnostica) for 30 min at room temperature (RT). Brain slices were then incubated for 20 h at RT with primary antibodies (chicken anti-GFP, 1:1000, ab13970, Abcam; rat anti-mcherry, 1:1000, Thermo Fisher Scientific, M11217) in PBS-based blocking solution containing 0.1% TX, 10% NGS and 0.001% sodium azide (S2002, Sigma-Aldrich). Brain slices were rinsed first in PBS-0.1% TX (2 x 10 min, RT), then in PBS (3 x 10 min, RT) and subsequently incubated with secondary antibodies (goat anti-chicken-FITC, Jackson #103-095-155, 1:500; goat anti-rabbit Alexa-Fluor-594, Thermo Fisher Scientific, A11012; 1:500) and DAPI (1:1000) for 2 hours at toom temperature. Brain slices were then rinsed in PBS-0.1% TX (2 x 10 min, RT) and PBS (3 x 10 min, RT), dehydrated in an ascending ethanol series, cleared with xylene (131769.1611, AppliChem), and mounted for imaging.

#### IHC in the pnVTA

Animals were transcardially perfused with 0.1 M phosphate-buffered saline (PBS) followed by 40 mL 4% paraformaldehyde (PFA). Brains were extracted and post-fixed in 4% PFA overnight and then transferred to 30% sucrose in PBS for cryoprotection. Brains were sectioned at 30 μm on a microtome and stored in a 0.1 M phosphate buffer at 4°C prior to immunohistochemistry. For behavioral cohorts, viral expression and optical-fiber placements were confirmed before inclusion in the presented datasets. Immunohistochemistry was performed as previously described. ^75,76^ In brief, free-floating sections were washed in 0.1 M PBS for 3 x 10 min intervals. Sections were then placed in blocking buffer (0.5% Triton X-100 and 5% natural goat serum in 0.1 M PBS) for 1 hour at room temperature. After blocking buffer, sections were placed in primary antibody (chicken anti-GFP, 1:2000, Abcam) overnight at 4°C. After 3 x 10 min 0.1 M PBS washes, sections were incubated in secondary antibody (AlexaFluor 488 goat anti-chicken, Abcam) for 2 hours at room temperature, followed by another round of washes (3 x 10 min in 0.1 M PBS, 3 x 10 min in 0.1 M PBS). After immunostaining, sections were mounted and coverslipped with Vectashield HardSet mounting medium containing DAPI (Vector Laboratories) and imaged on a Leica DM6 B microscope.

### Generation of OPRL1-Cre mouse line and reporter crosses

*Oprl1*^iresCre:GFP^ knock-in mice were generated at the University of Washington. A cassette encoding IRES-mnCre:GFP was inserted just 3’ of the termination codon in the last coding exon of the *Oprl1* gene. The 5′ arm (12 kb with *Pac*I and *Sal*I sites at 5’ and 3’ends, respectively) and 3′ arm (3.5 kb with *Xho*I and *Not*I sites at 5’and 3’ ends, respectively) of the *Oprl1* gene were amplified from a C57BL/6 BAC clone by PCR using Q5 Polymerase (New England Biolabs) and cloned into polylinkers of a targeting construct that contained IRES-mnCre:GFP, a frt-flanked Sv40Neo gene for positive selection, and HSV thymidine kinase and *Pgk*-diphtheria toxin A chain genes for negative selection. The IRES-mnCre:GFP cassette has an internal ribosome entry sequence (IRES), a myc-tag and nuclear localization signals at the N-terminus of Cre recombinase, which is fused to green fluorescent protein followed by a SV40 polyadenylation sequence (Cre:GFP). The construct was electroporated into G4 ES cells (C57Bl/6 × 129 Sv hybrid) and correct targeting was determined by Southern blot of DNA digested with *BamH*I using a 32P-labeled probe downstream of the 3′ arm of the targeting construct. Five of 77 clones analyzed were correctly targeted. One clone that was injected into blastocysts resulted in good chimeras that transmitted the targeted allele through the germline. Progeny were bred with *Gt(Rosa)26Sor*-FLP recombinase mice to remove the frt-flanked SV-Neo gene. Mice were then continuously backcrossed to C57Bl/6 mice.

To visualize OPRL1-Cre expression in the VTA, OPRL1-Cre mice were crossed to the Ai3 EYFP flox-stop reporter line (Jackson Lab, #007903). Adult OPRL1-Cre x Ai3 mice were transcardially perfused with 0.1 M phosphate-buffered saline (PBS) followed by 40 mL of 4% paraformaldehyde (PFA). Brains were extracted and post-fixed in 4% PFA overnight and then transferred to 30% sucrose in PBS for cryoprotection. Brains were sectioned at 30 μm on a microtome and stored in a 0.1 M phosphate buffer at 4°C. Midbrain sections were mounted on Super Frost Plus slides (ThermoFisher) and coverslipped with Vectashield HardSet mounting medium containing DAPI (Vector Laboratories) and imaged on a Leica DM6 B microscope.

### RNAscope fluorescent in situ hybridization (FISH)

Immediately after rapid decapitation of OPRL1-Cre mice (n = 2), brains were extracted, flash frozen in -50°C 2-methylbutane, and then stored at -80°C. Brains were sectioned coronally into 15 μm slices on a cryostat at -20°C, mounted onto Super Frost Plus slides (Fisher), and then stored at -80°C prior to RNAScope FISH. FISH was performed according to the RNAScope Fluorescent Multiplex Assay for use with fixed frozen tissues (Advanced Cell Diagnostics, Inc.). Slides with 2-4 brain sections each that contained the VTA were post-fixed in prechilled 10% neutral-buffered formalin for 15 minutes at 4°C, dehydrated in ethanol, and then treated with a protease IV solution for 30 minutes at 40°C. The sections were then incubated for 2 hours at 40°C with target probes (Advance Cell Diagnostics, Inc.) for mouse *Oprl1* (accession number NM_011012.5, target region 988 – 1937), *Th* (accession number NM_009377.1, target region 483 – 1603), and Cre (target region 2 – 972). Next, sections underwent a series of probe amplifications (AMP1-4) at 40°C, including a final incubation with fluorescently labelled probes (Alex 488, Atto 550, Atto 647) targeted to the specified channels (C1-C3) that were associated with each of the probes. Finally, sections were stained with DAPI and slides were coverslipped with Vectashield Hard-Set mounting medium (Vector Laboratories). Slides were imaged on a Leica TCS SPE confocal microscope (Leica) at 60x magnification, and Fiji and HALO software were used to process images and quantify expression. Images were obtained under consistent threshold and exposure time standards. Leica images were opened and converted to TIFs for compatibility with HALO software. In HALO, cell ROIs were first made using the DAPI channel, and then *Oprl1*+ and Cre+ cells were identified if a fluorescent threshold for each channel was met within the cell ROI. 2-3 separate slices were quantified for each animal.

### Photometry Recording

#### Recordings in ARC

For fibre photometry studies in the arcuate nucleus, the set-up of the photometry recorder ^77^ consisted of an RZ5P real-time processor (Tucker-Davis Technologies) connected to a light source driver (LED Driver; Doric Lenses). The LED Driver constantly delivered excitation light at 405 nm (control) and 465 nm (NOPLight) wavelengths. The light sources were filtered by a four-port fluorescence minicube (FMC_AE(405)_E1(460-490)_F1(500-550)_S, Doric Lenses) before reaching the animal. The fluorescence signals were collected from the same fibre using a photoreceiver (Model 2151, New Focus), sent back to the RZ5P processor, and gathered by Synapse software (v.95-43718P, Tucker-Davis Technologies).

#### Recordings in pnVTA

For fiber-photometry studies in the pnVTA, recordings were made continuously throughout the entirety of the pharmacology (30 min), chemogenetic (55 min), head-fixed sucrose (25 min), tail lift (20 min), and conditioned reward-seeking (60 min) sessions. Prior to recording, an optic fibre was attached to the implanted fibre using a ferrule sleeve (Doric, ZR_2.5). In pharmacology and conditioned reward-seeking experiments, a 531-Hz sinusoidal LED light (Thorlabs, LED light: M470F3; LED driver: DC4104) was bandpass filtered (470 ± 20 nm, Doric, FMC4) to excite NOPLight and evoke NOPR-agonist dependent emission while a 211-Hz sinusoidal LED light (Thorlabs, M405FP1; LED driver: DC4104) was bandpass filtered (405 ± 10 nm, Doric, FMC4) to excite NOPLight and evoke NOPR-agonist independent isosbestic control emission. In chemogenetics, head-fixed sucrose, and tail lift experiments, a 531-Hz sinusoidal LED light (Thorlabs, LED light: M490F3; LED driver: DC4104) was bandpass filtered (490 ± 20 nm, Doric, FMC6) to excite NOPLight and evoke NOPR-agonist dependent emission while a 211-Hz sinusoidal LED light (Doric, CLED_435; Thorlabs, LED driver: DC4104) was bandpass filtered (435 ± 10 nm, Doric, FMC6) to excite NOPLight and evoke NOPR-agonist independent isosbestic control emission. Prior to recording, a minimum 120s period of NOPLight excitation with either 470-nm and 405-nm or 490-nm and 435-nm light was used to remove the majority of baseline drift. Power output for each LED was measured at the tip of the optic fibre and adjusted to 30 μW before each day of recording. NOPLight fluorescence traveled back through the same optic fibre before being bandpass filtered (525 ± 25 nm, Doric, FMC4 or FMC6), detected with a photodetector system (Doric, DFD_FOA_FC), and recorded by a real-time processor (TDT, RZ5P). The 531-Hz and 211-Hz signals were extracted in real time by the TDT program Synapse at a sampling rate of 1017.25 Hz.

### In vivo animal experiments

All animal behaviors were performed within a sound-attenuated room maintained at 23°C at least one week after habituation to the holding room. Animals were handled for a minimum of three days prior to experimentation, as well as habituated to the attachment of a fibre photometry patch cord to their fibre implants. For all experiments, mice were brought into the experimental room and allowed to acclimate to the space for at least 30 min prior to beginning any testing. All experiments were conducted in red light to accommodate the reverse light cycle schedule, unless otherwise stated. All pharmacological interventions were administered in a counterbalanced manner. All sessions were video recorded.

#### In vivo pharmacology experiments in the ARC

Mice injected with AAV-DJ-hSyn-NOPLight were acclimatized to the behavior set-up 3 weeks post-surgery for one week. Awake animals were placed individually in a box and an optic fibre cable was connected to the implanted fibre. The optic fibre was attached to a swivel joint above the box to avoid moving limitations. Recording started 5 min after optic fibre tethering. A 10 min-long baseline was recorded prior to the i.p. injection of vehicle or RO 64-6198 (v doses). Animals have no access to water or food during the recording session.

#### In vivo pharmacology experiments in the pnVTA

WT mice injected with either AAV-DJ-hSyn-NOPLight (n = 16) or AAV-DJ-hSyn-NOPLight-ctr (n = 6) and OPRL1*-*Cre mice injected with AAV-DJ-hSyn-FLEX-NOPLight (n = 3) were allowed to recover a minimum of 3 weeks after surgery. Three days before testing they were habituated to handling, fibre photometry cable attachment, and to the behavioral test box. On test day, animals were placed into the behavioral test box, which was a 10” x 10” clear acrylic box with a layer of bedding on the floor illuminated by a dim, diffuse white light (30 lux). Fibre photometry recordings were made using a 405 nm LED as the isosbestic channel and a 470 nm LED as the signal channel. After starting the photometry recording, the mice were free to move around the box with no intervention for 5 min to establish a baseline photometry signal. At 5 min into the recording, NOPLight mice (n = 16) were scruffed and received an intraperitoneal (i.p.) injection of vehicle or 1, 5, or 10 mg/kg of the selective NOPR agonist Ro 64-6198 and were recorded for an additional 25 min. NOPLight-ctr and FLEX-NOPLight mice received an i.p. injection of 10 mg/kg Ro 64-6198 5 min into the recording. Two subsets of the NOPLight animals were recorded on three separate, counterbalanced days (at least 24 hours apart). The first subset (n = 4) received either i) an i.p. injection of 10 mg/kg Ro 64-6198 5 min into the recording (RO), ii) an oral gavage (o.g.) treatment with 10 mg/kg of selective NOPR antagonist LY2940094 5 min into the recording (LY), or iii) an o.g. treatment with 10 mg/kg LY2940094 30 min prior to the recording followed by an i.p. injection with 10 mg/kg Ro 64-6198 5 min into the recording (LY pretreatment + Ro). The second subset (n = 4-9) received either i) an i.p. injection of 10 mg/kg Ro 64-6198 5 min into the recording (RO), ii) an i.p. injection of 10 mg/kg of selective NOPR antagonist J-113397 5 min into the recording (J11), or iii) an i.p. injection of 10 mg/kg J-113397 30 min prior to the recording followed by an i.p. injection with 10 mg/kg Ro 64-6198 5 min into the recording (J11 pretreatment + Ro).

#### In vivo chemogenetics (DREADD) experiments

PNOC*-*Cre mice co-injected with AAVDJ-hSyn-FLEX-NOPLight and either AAV5-EF1a-DIO-HA-hM3D(Gq)-mScarlet (n = 8) or AAV5-DIO-hSyn-mCherry (n = 3) were allowed to recover a minimum of 3 weeks after surgery. Three days before testing they were habituated to handling, fibre photometry cable attachment, and to the behavioral test box. On test day, animals were placed into the behavioral test box, which was a 10” x 10” clear acrylic box with a layer of bedding on the floor illuminated by a dim, diffuse white light (30 lux). Fibre photometry recordings were made using a 435 nm LED as the isosbestic channel and a 490 nm LED as the signal channel. After starting the photometry recording, the mice were free to move around the box with no intervention for 10 min to establish a baseline photometry signal. At 10 min into the recording, mice were scruffed and received an intraperitoneal (i.p.) injection of 5 mg/kg clozapine-N-oxide (CNO) and were recorded for an additional 45 min. A subset of the DIO hM3D(Gq) animals (n = 3) were recorded on two separate, counterbalanced days (at least 24 hours apart). On one recording day, they received the CNO treatment and recording timeline described above. On the other recording day, they were administered an oral gavage (o.g.) treatment with 10 mg/kg of the selective NOPR antagonist LY2940094 30 min prior to the photometry recording. The LY-pretreatment group then underwent the same recording timeline and CNO injection (5 mg/kg i.p., after a 10 min baseline) as the CNO only day.

#### Head-fixed cued sucrose access paradigm

WT mice injected with AAV-DJ-hSyn-NOPLight (n = 7) and implanted with an optic fibre and stainless-steel head-ring to allow for head-fixation during fibre photometry recording were allowed to recover a minimum of 3 weeks after surgery. One week prior to behavioral testing, mice were food restricted down to 90% of their free feeding body weight. For the four days prior to testing, animals were habituated to handling, fibre photometry cable attachment, and head-fixation. Animals were head-fixed to minimize motion-related artefacts in the fibre photometry signal, and all head-fixed testing was completed using the open-source OHRBETS platform. ^78^

Fibre photometry recordings of NOPLight signal during tone-cued access to 10% sucrose solution were made over two counterbalanced sessions where animals received an i.p. injection 30 min prior to the recording of either i) 20 mg/kg NOPR antagonist J-113397 or ii) vehicle. Each session consisted of a 5-minute baseline period where animals were head-fixed with no stimuli delivery, 15 tone-cued sucrose trials, and then a 5-minute baseline period at the end of the session (25 minutes total). Each cued sucrose trial consisted of a 5-second auditory tone (4kHz, 80dB) immediately followed by a 5-second extension of a retractable lick spout and delivery of five pulses of 10% sucrose solution (1.5 μL/pulse, 200ms inter-pulse interval) where mice could lick the spout to consume the solution. Cued sucrose trials were separated by a variable inter-trial interval of 45 to 75 seconds.

All behavioral hardware were controlled using an Arduino Mega 2560 REV3 (Arduino) and custom Arduino programs. Individual licks were detected using a capacitive touch sensor (Adafruit MPR121) that was attached to the retractable lick spout. The pulsed sucrose delivery was controlled by a solenoid (Parker 003-0257-900). The timing of solenoid openings and lick events were recorded and synchronized with the photometry signal via TTL communication from the Arduino Mega to the fibre photometry system. Fibre photometry recordings were made using a 435 nm LED as the isosbestic channel and a 490 nm LED as the signal channel.

#### Tail lift behavioral experiments

PNOC*-*Cre mice injected with AAV-DJ-EF1a-DIO-GCaMP6m (n = 4), OPRL1*-*Cre mice injected with AAV-DJ-hSyn-FLEX-NOPLight (n = 3), and WT mice injected with AAV-DJ-hSyn-NOPLight-ctr (n = 3) were allowed to recover a minimum of 3 weeks after surgery. Three days before testing they were habituated to handling, fibre photometry cable attachment, and to the behavioral test box. On test day, animals were placed into the behavioral test box, which was a 10” x 10” clear acrylic box illuminated by a dim, diffuse white light (30 lux). Fibre photometry recordings were made using a 435-nm LED as the isosbestic channel and a 490-nm LED as the signal channel. After starting the photometry recording, mice were free to move around the box with no intervention for 5 minutes to establish a baseline photometry signal. After the 5-minute baseline, mice underwent four tail lift trials where they were suspended by the tail for 10 seconds and then gently returned to the behavioral test box. Tail lift trials were separated by a variable inter-trial interval of 120-300 seconds. All suspensions were made to the same height. After the final trial, photometry recording continued for an additional 5 minutes to establish a post-test signal baseline.

#### Reward-seeking (Pavlovian, operant) conditioning paradigms

One week prior to Pavlovian conditioning and 3 weeks after surgery, WT fibre photometry mice expressing AAV-DJ-hSyn-NOPLight in the VTA (n = 4) were food restricted down to 90% of their free feeding body weight. All reward-seeking training was completed in Med-Associates operant conditioning boxes (ENV-307A). Fibre photometry recordings were made using a 405nm LED as the isosbestic channel and a 470 nm LED as the signal channel. Mice were first trained to associate illumination of a house light (CS, 5 s) with delivery of a single sucrose pellet (US) occurring immediately after the house light turned off. A randomized intertrial interval of between 30-120 s separated consecutive trials. Pavlovian conditioning sessions lasted for 60 min, duirng which an average of 36-38 rewards were presented. Pavlovian conditioning was repeated over 5 days a total of 5 times, with simultaneous fibre-photometry recordings made during session 1 and session 4. Animals were then moved onto a fixed ratio 1 (FR1) schedule for 5 days (60 min/session), where they were required to perform a nosepoke in the active nosepoke port one time to receive the 5 s house light cue and subsequent pellet delivery. Pokes made into the inactive port had no effect. Simultaneous fibre-photometry recordings were made during FR1 sessions 1 and 4. Following FR1 training, the ratio was increased to an FR3 schedule for 4 days (60 min/session) requiring the mice to perform 3 active port nosepokes to receive the house light cue and a sucrose pellet, with simultaneous photometry recording during session 3. Last, mice were placed in a single, 120 min session on a progressive ratio schedule (PR) where the nosepoke criteria for each subsequent reward delivery followed the geometric progression *n*_*i*_ *= 5e*^*i/5*^ *– 5* (1, 2, 4, 6, 9, 12…), increasing in an exponential manner.

### Data Analysis for Photometry Recordings

#### Recordings in ARC

Fibre-photometry data was pre-processed by down sampling the raw data to 1 Hz and removing the first and last seconds of each recording to avoid noise. To correct for bleaching, the fluorescence decay as the baseline (F_0_) for both signal and control channel were fitted using a Power-like Model^47^. If no model could be fitted for a sample (e.g. no decay), median of the baseline recording was used as a substitute. The relative change post-injection (ΔF/F_0_) was estimated by

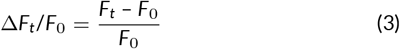

for both the signal and control channel separately (where F_t_ is the raw signal at time *t*). ΔF/F_0control_ was subsequently subtracted from ΔF/F_0signal_ to correct for motion artifacts, obtaining a final estimate of the relative change in fluorescence intensity for each sample.

#### Recordings in pnVTA

Custom MATLAB scripts were developed for analyzing fibre-photometry data in the context of mouse behavior. A linear least squares (LLS) fit was applied to the control signal (405 or 435 nm) to align and fit it to the excitation signal (470 or 490 nm). For pharmacology and chemogenetics experiments where the entire length of the recording was analyzed to evaluate long-term changes in NOPLight fluorescence following drug injection, LLS fit was calculated using the recording’s ‘baseline’ period that preceded the injection (5-10 min) and the fitted isosbestic signal was subtracted from the excitation signal to detrend bleaching and remove movement artefacts. To reduce high-frequency noise, data were down-sampled by a factor of 300. The processed fibre photometry trace was then smoothed across a rolling 10 s window, and z-scored relative to the mean and standard deviation of the baseline period preceding drug injection (first 5 or 10 min of the recording for pharmacology and chemogenetics experiments, respectively).

For all behavioral experiments (head-fixed sucrose, tail lift, conditioned reward seeking) where short epochs were evaluated, decay from bleaching was first detrended by fitting a 4^th^ degree polynomial function to the raw signal and isosbestic traces, then dividing by the resulting curve. Next, the LLS fit of the isosbestic signal to the excitation signal was calculated over the entire session and the excitation signal was normalized by dividing the resulting fitted isosbestic signal. To reduce high-frequency noise, data were down-sampled by a factor of 100. The processed traces were then smoothed across a rolling 1 s window, extracted in windows surrounding the onset of relevant behavioral events (e.g., nose poke, cue onset, reward delivery, tail lift), z-scored relative to the mean and standard deviation of each event window, and then averaged. The post-processed fibre photometry signal was analyzed in the context of animal behavior during Pavlovian conditioning and operant task performance.

### Statistical analyses

All data were averaged and expressed as mean ± SEM. Statistical significance was taken as *p < 0.05, ** p < 0.01, ***p < 0.001, and ****p < 0.0001, as determined by Mann-Whitney test, Wilcoxon test, two-way repeated-measures ANOVA followed by Bonferroni post hoc tests as appropriate. All n values for each experimental group are described in the corresponding figure legend. For behavioral experiments, group size ranged from n = 3 to n = 16. Statistical analyses were performed in GraphPad Prism 9 (GraphPad, La Jolla, CA) and MATLAB 9.9 (The MathWorks, Natick, MA).

