## Supplementary Figure 1 - 10 for "Development of a genetically encoded sensor for probing endogenous nociceptin opioid peptide release"

### SUPPLEMENTARY FIGURES

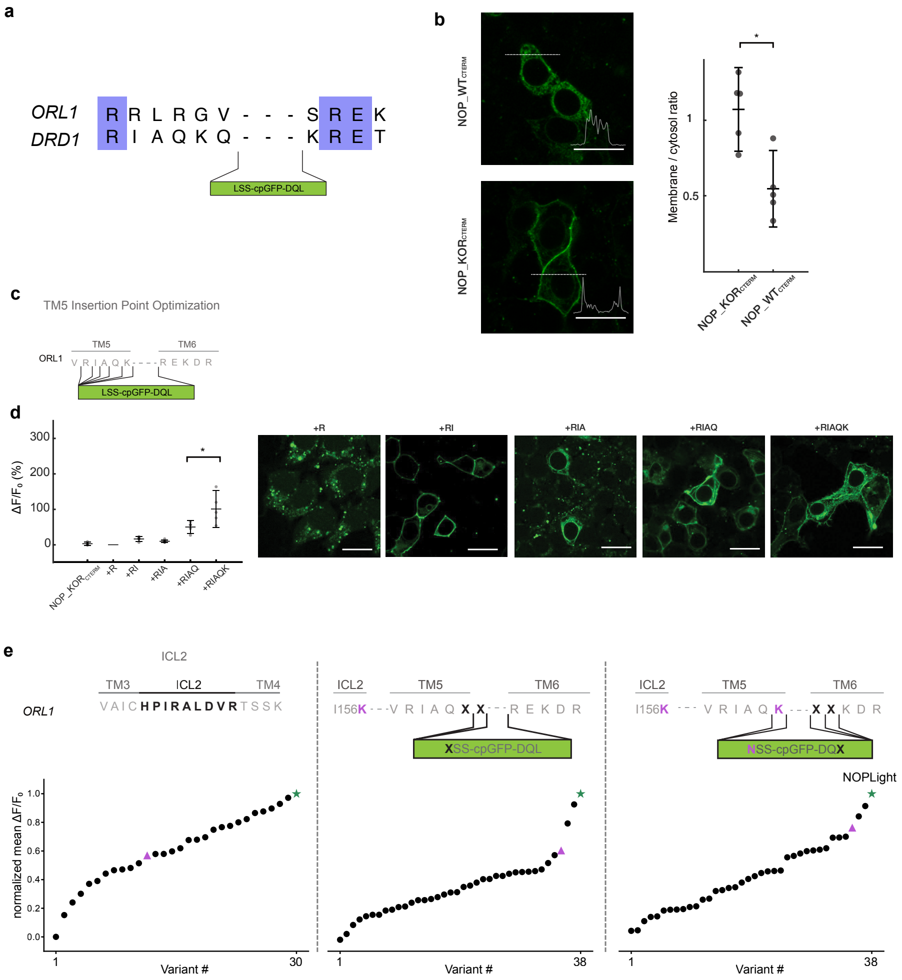


**SUPPLEMENTARY FIGURE 1** **Development and optimization of NOPLight.**

**a.** Sequence alignment of transmembrane domains (TM) 5 and 6 from OPRL1 and DRD1 to determine the insertion site for the cpGFP module from dLight1.3. Color code indicates percent identity between the human NOPR and DRD1. **b.** Left: Representative images of membrane expression profile of HEK293T cells expressing *in silico* designed sensor prototype with a wild-type NOPR C-terminus (N/OFQ$_{_WTcterm}$) and a chimeric sensor with the C-terminus of kappa opioid receptor (N/OFQ$_{_KORcterm}$). Insets: fluorescent intensity of all pixels along the dotted line of each image. Scale bars, 10 µm. Right: Quantification of the ratio between pixel averaged fluorescent intensity of cell membrane and that of the cytosol. n = 5 cells. [illegible]P = 0.011, one-sided Mann Whitney U test[illegible] **c.** Schematic representation of the optimization of the insertion site at TM5 linker region. **d.** Left: Maximal $\Delta F/F_0$ response to 10 µM N/OFQ of all sensor prototypes shown in **c.** Right: representative image of HEK293T cells expressing sensor prototypes. **e.** Top: Schematic representation of three rounds of directed mutagenesis to improve sensor dynamic range. Amino acids that were mutated into a subset of different amino acids in the screening process are labelled in bold. Selected mutation from previous round of screening labelled in magenta. Bottom: Normalized $\Delta F/F_0$ response of HEK293T cells expressing each variant to 10 µM N/OFQ. Each data point represents averaged $\Delta F/F_0$ response from > 3 cells in 3 independent experiments normalized to the maximal $\Delta F/F_0$ response of the best variant in the current round of screening. Magenta: Selected variant from previous round of screening. Green: Best variant in current round of screening.

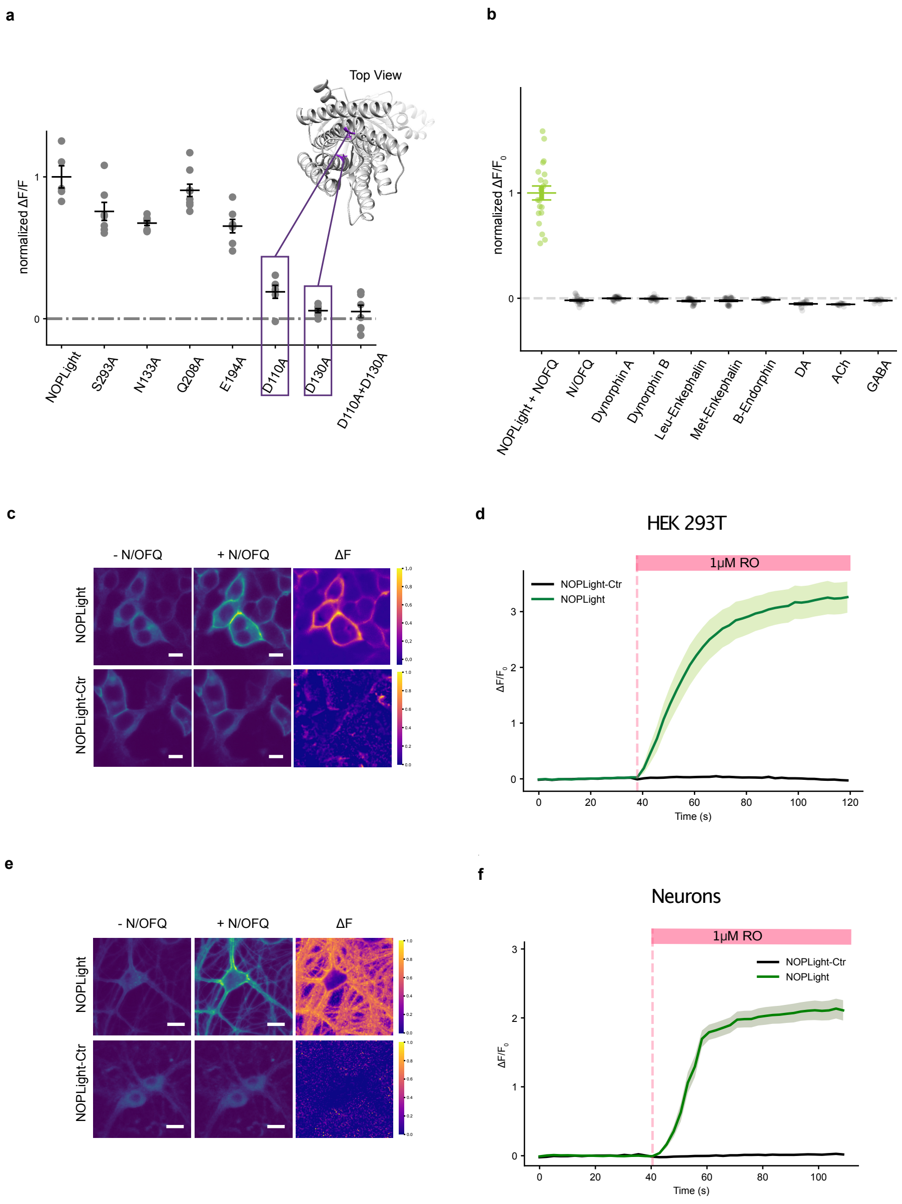


**SUPPLEMENTARY FIGURE 2 Development and characterisation of NOPLight-ctr.**
**a.** Quantification of maximal $\Delta F/F_0$ in response to 10 µM N/OFQ in HEK 293T cells expressing NOPLight mutants. Mutated residues indicated by their absolute residue numbering in reference to human NOPR. Structural model of NOPLight-ctr is predicted by RoseTTAfold[42] The two mutations selected for NOPLight-ctr are highlighted in purple. **b.** Normalized maximal $\Delta F/F_0$ of NOPLight-Ctr-expressing HEK 293T cells in response to endogenous opioid peptides (1 µM) and fast neurotransmitters (DA: dopamine, Ach: acetylcholine, GABA: gamma-Aminobutyric acid (1 mM)). All data normalized to NOPLight-expressing HEK293T cells (green). **c.** Representative images of HEK 293T cells expressing NOPLight (top; scale bars, 10 µm) and NOPLight-ctr (bottom; scale bars, 10 µm) before and after application of 1 µM Ro 64-6198. Corresponding normalized pixelwise ΔF shown on the right. **d.** Average fluorescent-fold change ($\Delta F/F_0$) of HEK 293T cell expressing NOPLight (light green trace) or NOPLight-ctr (black trace) in response to 1 µM RO 64-6198 (3 experiments, data shown as mean + s.e.m). **e.** Representative images of neurons expressing NOPLight (top; scale bars, 20 µm) and NOPLight-ctr (bottom; scale bars, 20 µm) before and after application of 1 µM Ro 64-6198. Corresponding normalized pixelwise ΔF shown on the right. **f.** Average fluorescent-fold change ($\Delta F/F_0$) of neurons expressing NOPLight (light green trace) or NOPLight-ctr (black trace) in response to 1 µM RO 64-6198 (3 experiments, data shown as mean ± s.e.m).

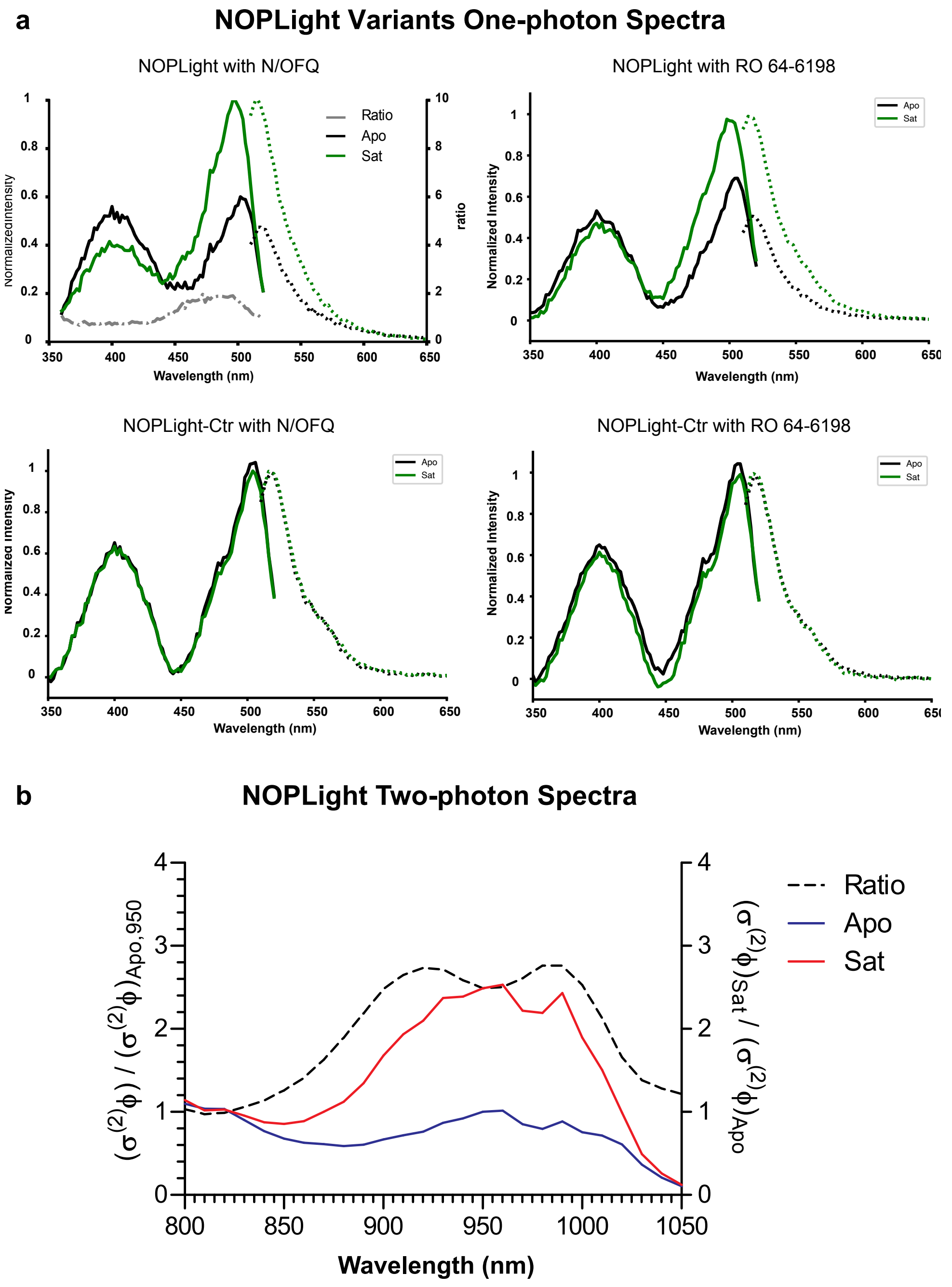


**SUPPLEMENTARY FIGURE 3 Development and characterisation of NOPLight-ctr.**

**a.** Top, normalized one-photon fluorescence excitation (solid lines, $\lambda_{excitation}$ = 360 – 520 nm, $\lambda_{emission}$ = 560 nm) and emission (dotted lines, $\lambda_{excitation}$ = 470 nm, $\lambda_{emission}$ = 510 – 650 nm) spectra of NOPLight-expressing HEK293T cells in the absence and presence of either N/OFQ (1 µM) or RO 64-6198 (1 µM). Traces shown are averaged from 3 independent experiments. Intensity measured at each wavelength is normalized to the maximum intensities measured for both the excitation and emission spectra. Bottom, same as on top with NOPLight-Ctr-expressing HEK 293T cells in the absence and presence of N/OFQ (1 µM) or RO 64-6198 (1 µM). **b.** Relative two-photon brightness of NOPLight imaged in transfected HEK cells grown attached to a glass coverslip in the presence (Sat) or absence (Apo) of N/OFQ (1 µM). Ratio between Sat and Apo shown in dashed line. Each trace is the average of 3 independent experiments.

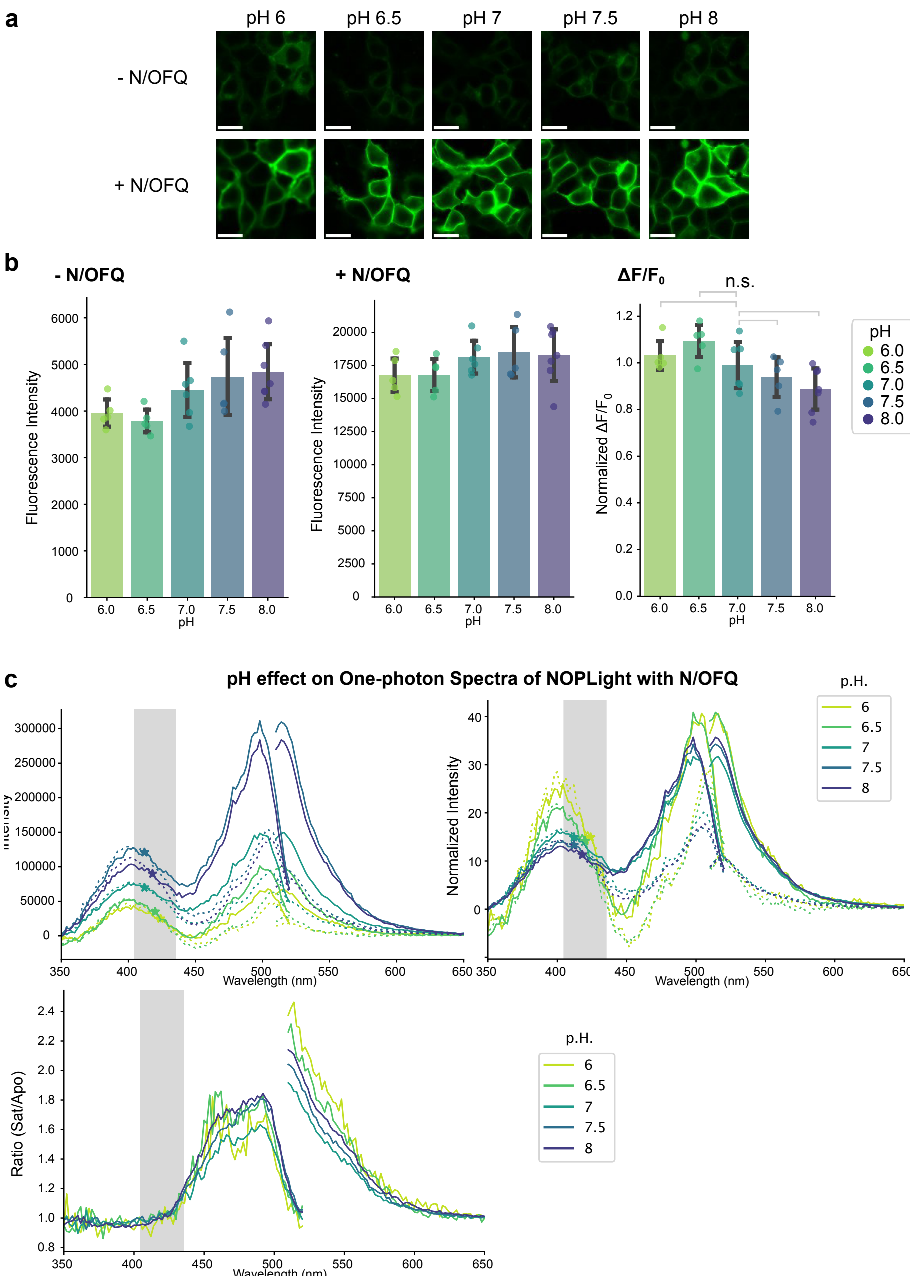


**SUPPLEMENTARY FIGURE 4 Characterization of NOPLight sensitivity to external p.H.**

**a.** Representative images of NOPLight-expressing HEK 293T cells in the absence (top) and presence (bottom) of 1 µM N/OFQ. Scale bar: 20 µm. **b.** Quantification of fluorescence intensity in the absence (left) and presence (middle) of 1 µM N/OFQ, and fluorescent-fold change (ΔF/F) normalized to the ΔF/F measured at extracellular p.H. 7 (right, ANOVA with Tukey Kramer post-hoc test of all p.H. conditions compared to p.H, 7. P = 0. 942, 0.358, 0.883 and 0.289 respectively). Data shown as mean ± SEM. **c.** Top left: One-photon spectra of NOPLight-expressing HEK 293T cells at various extracellular p.H. in the presence (Sat, solid lines) and absence (Apo, dashed lines) of N/OFQ (1 µM). Isosbestic point at each p.H. marked with " * ". Top right: One-photon spectra of NOPLight-expressing HEK 293T cells at various p.H. normalized by the corresponding area under the curve of with the presence of N/OFQ at each extracellular p.H. condition. Bottom: Ratio between Sat and Apo at various extracellular p.H.. Shaded bar indicates the range between 405 to 435 nm.

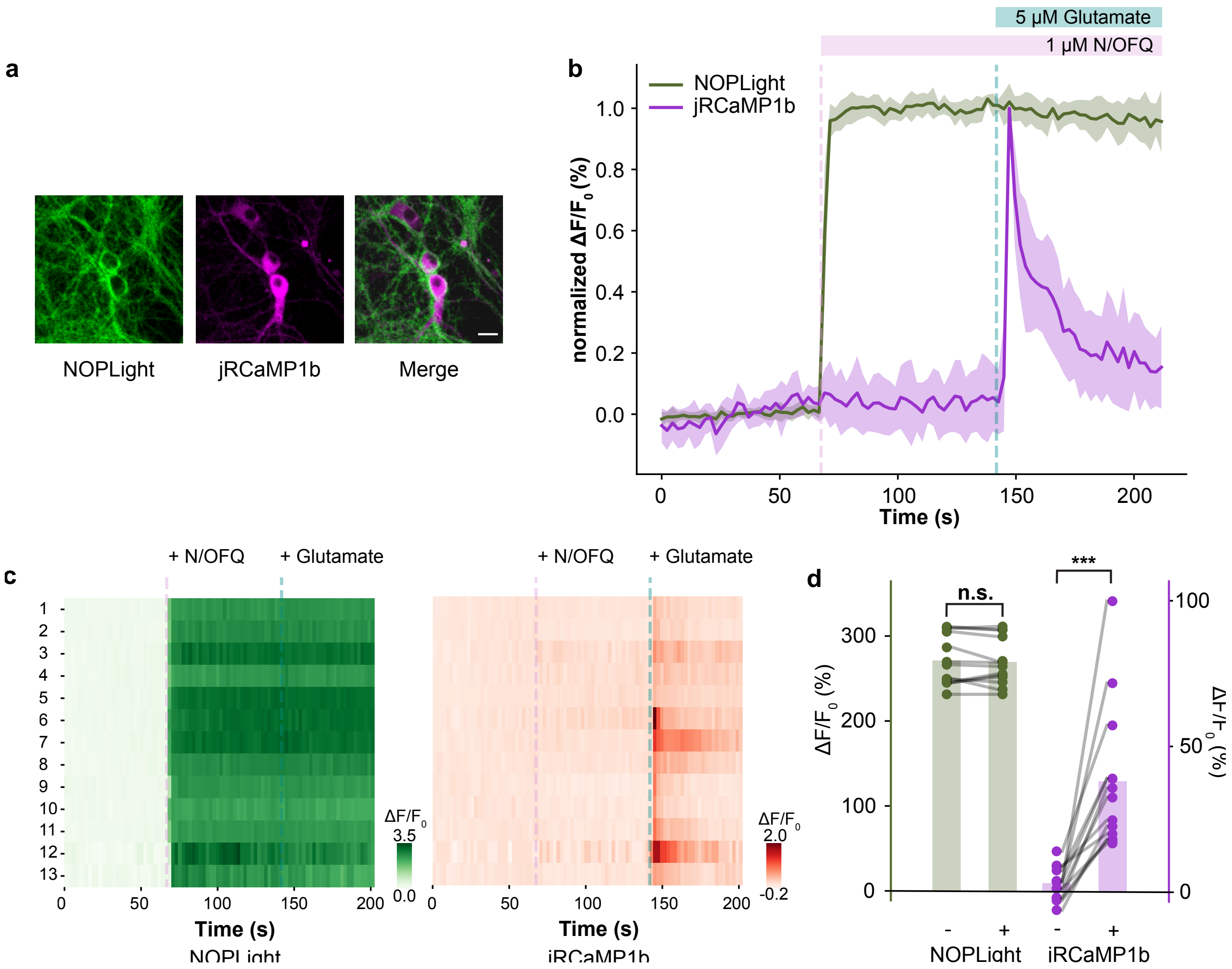


**SUPPLEMENTARY FIGURE 5 Changes in neuronal activity do not affect the fluorescence response of NOPLight to N/OFQ.**
**a.** Representative image of neurons co-expressing NOPLight (right) and jRCaMP1b (middle). Scale bar: 20 μm. **b.** Normalized average fluorescent-fold change ($\Delta F/F_0$) of NOPLight (green) and jRCaMP1b (magenta) in response to 1 μM N/OFQ followed by 5 μM glutamate. (13 neurons from 2 independent experiments, data shown as mean ± s.t.d). **c.** Corresponding heat map of **b**, where each row corresponds to one neuron. Left: NOPLight fluorescent response; Right: jRCaMP1b fluorescent response. **d.** Quantification of average $\Delta F/F_0$ (%) from **b**. NOPLight (left Y-axis, green) and jRCaMP1b (right Y-axis, magenta) signals were quantified 30 seconds before (-) and after (+) application of glutamate (paired sample t-test, ***$p<0.001$).

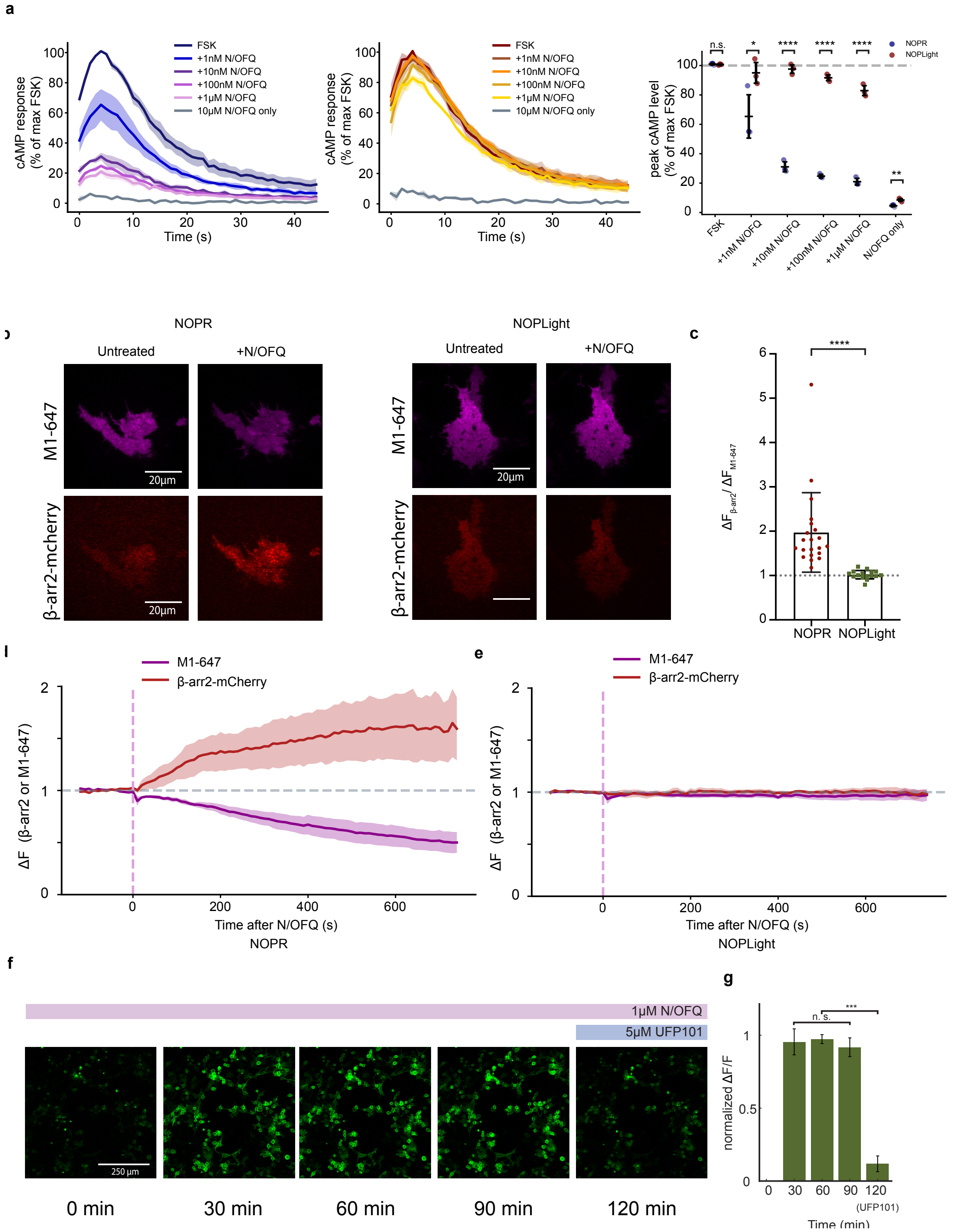


**SUPPLEMENTARY FIGURE 6** Characterization of sensor coupling to intracellular signaling pathways.

**SUPPLEMENTARY FIGURE 6 Characterization of sensor coupling to intracellular signaling pathways.**

**a**. Left: cAMP GloSensor luciferase activity traces in NOPR expressing HEK293 cells in response to different concentrations of N/OFQ, normalized to the peak activity evoked by 3µM of Forskolin (FSK). Middle: cAMP GloSensor luciferase activity traces in NOPLight expressing HEK293 cells in response of different concentrations of N/OFQ, normalized to the peak activity evoked by 3 µM of Forskolin (FSK). Right: Quantification of peak cAMP activity of NOPR and NOPLight expressing cells at different concentrations of N/OFQ. (n = 3 independent experiments, data shown as mean ± SEM, two sample t-test, n.s. P = 0.584, *P = 0.0623, **P = 0.00874, ****P < 0.0001). **b**. Representative images of cells co-expressing mCherry-tagged β-arrestin-2 with NOPR (left, scale bar = 20 µm) or NOPLight (right, scale bar = 20 µm) before and 12.5 min after N/OFQ (10 µM) stimulation. NOPR and NOPLight are labeled with Alexa-647-conjugated M1 anti-FLAG antibody (M1-647). **c**. Ratio of change in mCherry-tagged β-arrestin-2 signal versus change of M1-647 signal in NOPR and NOPLight expressing cells. (n = 21 and 16 cells respectively, from 3 independent experiments, data shown as mean ± s.d., one-sided Mann-Whitney U test, P<0.0001). **d**. Plasma membrane signal (TIRF) of mCherry-tagged β-arrestin-2 and FLAG-tagged NOPR signal before and after the application of N/OFQ. (n = 3 independent experiments, data shown as mean ± s.e.m) **e.** Similar to **d**, cells expressing NOPLight instead of NOPR. **f.** representative images of NOPLight expressing HEK293T cells at 0 min, 30 min, 60 min, 90 min and 120 min after the addition of N/OFQ (1µM). 5 µM UFP 101 was added at t = 110 min. **g.** Normalized fluorescent response ($\Delta F/F_0$) at time points as in **f.** (n = 3 independent experiments, P = 0.7522 for one-way repeated measures ANOVA test between fluorescence response at 30, 60 and 120 min after N/OFQ, P<0.001 for one-way repeated measures ANOVA test between 30, 60, 120 min after N/OFQ and after UFP 101.).

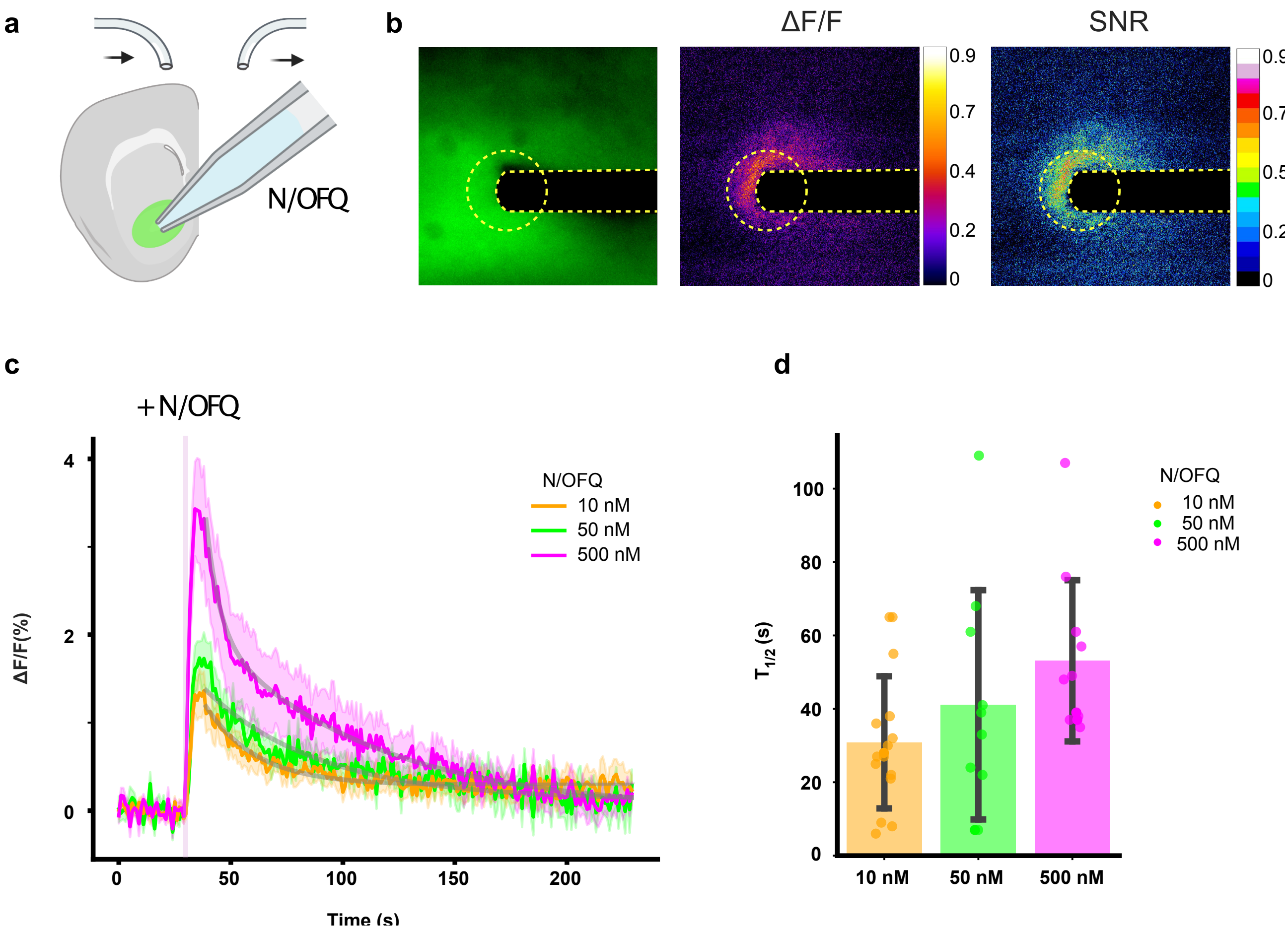


**SUPPLEMENTARY FIGURE 7 NOPlight kinetic measurements in brain slices.**

**a.** Schematic representation of AAV-DJ-hSynapsin1-NOPLight injection into the NAc. **b.** Left: Representative image of acute brain slice expressing NOPLight; middle: pixel-wise ΔF/F immediately after application of N/OFQ; right: pixel-wise signal to noise ratio immediately after application of N/OFQ. **c.** Time-course plot of ΔF/F traces depicting the response under each concentration of N/OFQ. Data shown as mean (solid line) ± SEM (shaded area). Gray traces represent fitted mono-exponential decay model. Shaded pink bar represents the time point at which N/OFQ was applied. n = 16, 10, 11 slices respectively with 10, 50 and 500 nM N/OFQ from N = 5 animals. **d.** Quantification of the decay half-time at different concentrations of N/OFQ. n = 16, 10, 11 slices respectively. Bar plot represents mean ± SEM.

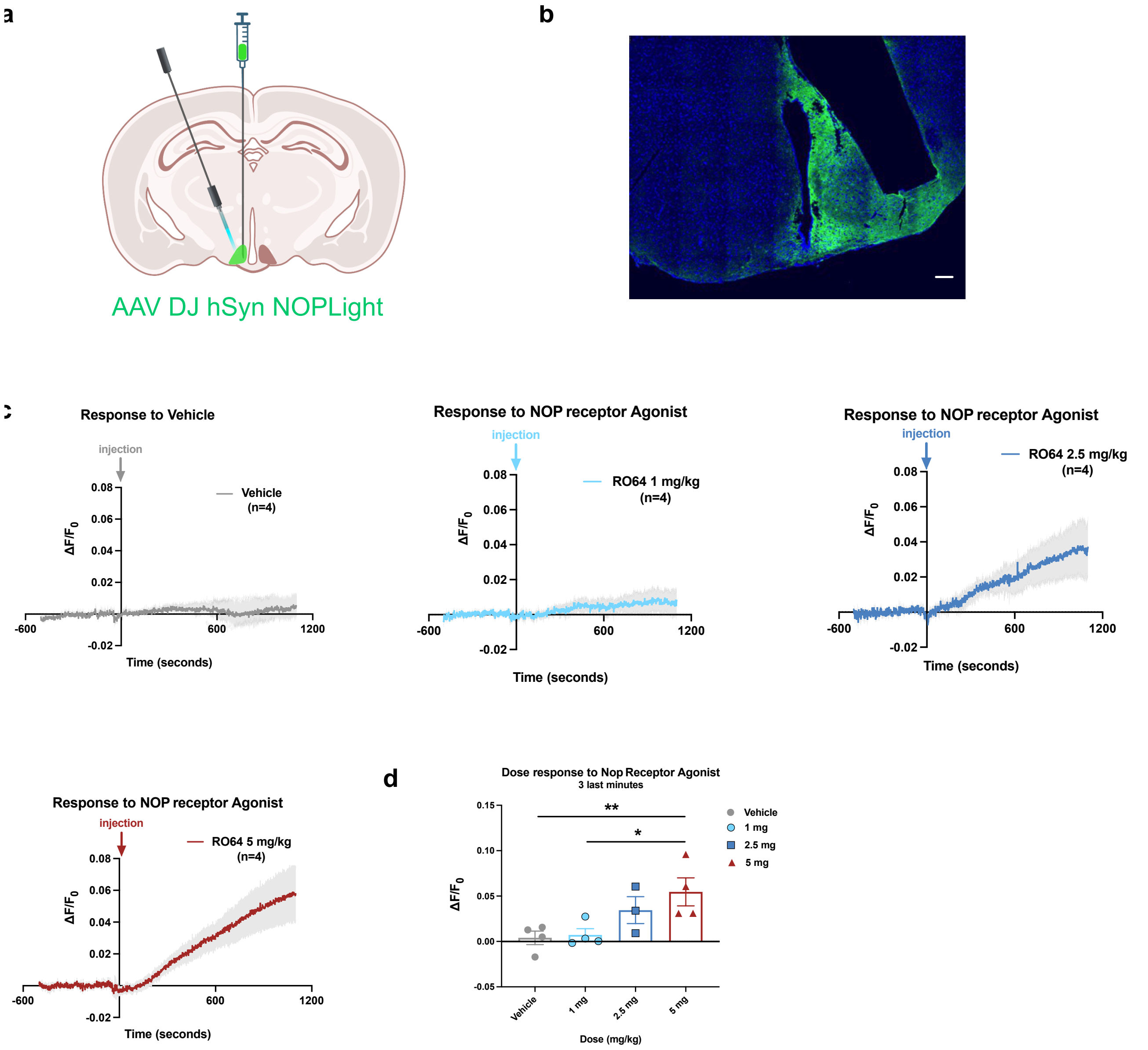


**SUPPLEMENTARY FIGURE 8 Target engagement of a NOPR agonist in the arcuate nucleus.**
**a.** Schematic representation showing AAV-DJ-hSynapsin1-NOPLight (NOPsyn) (0.8 x 1013 GC/mL) injection into the arcuate nucleus of the hypothalamus (ARC), followed by optic fibre implantation. **b.** Representative image of NOPLight-expressing brain slice showing optic fibre placement. GFP (green) expression, and DAPI in the ARC. Scale bar, 100 μm. **c**. Average *in vivo* photometry trace ($\Delta F/F_0$) for vehicle response. **d**. Average fluorescence traces showing the response to RO64 injection at 1 mg/kg. **e**. Average fluorescence traces showing the response to RO64 at 2.5 mg/kg. **f**. Average fluorescence traces showing the response to RO64 at 5 mg/kg. All injections were performed I.P. **g**. Quantification of average fluorescence signals in 3-min bins (957-1137 s) at the end of fibre photometry recordings, including data from (**c-f**) vehicle response and the response to RO64 (1, 2.5, and 5 mg/kg) IP injection. n=5 mice. Data are expressed as mean ± standard error of the mean (SEM). One-way ANOVA with Tukey's multiple comparisons test. Vehicle vs. RO64 5 mg/kg **P= 0.0058. RO64 1 mg/kg vs. RO64 5 mg/kg *P= 0.0131.

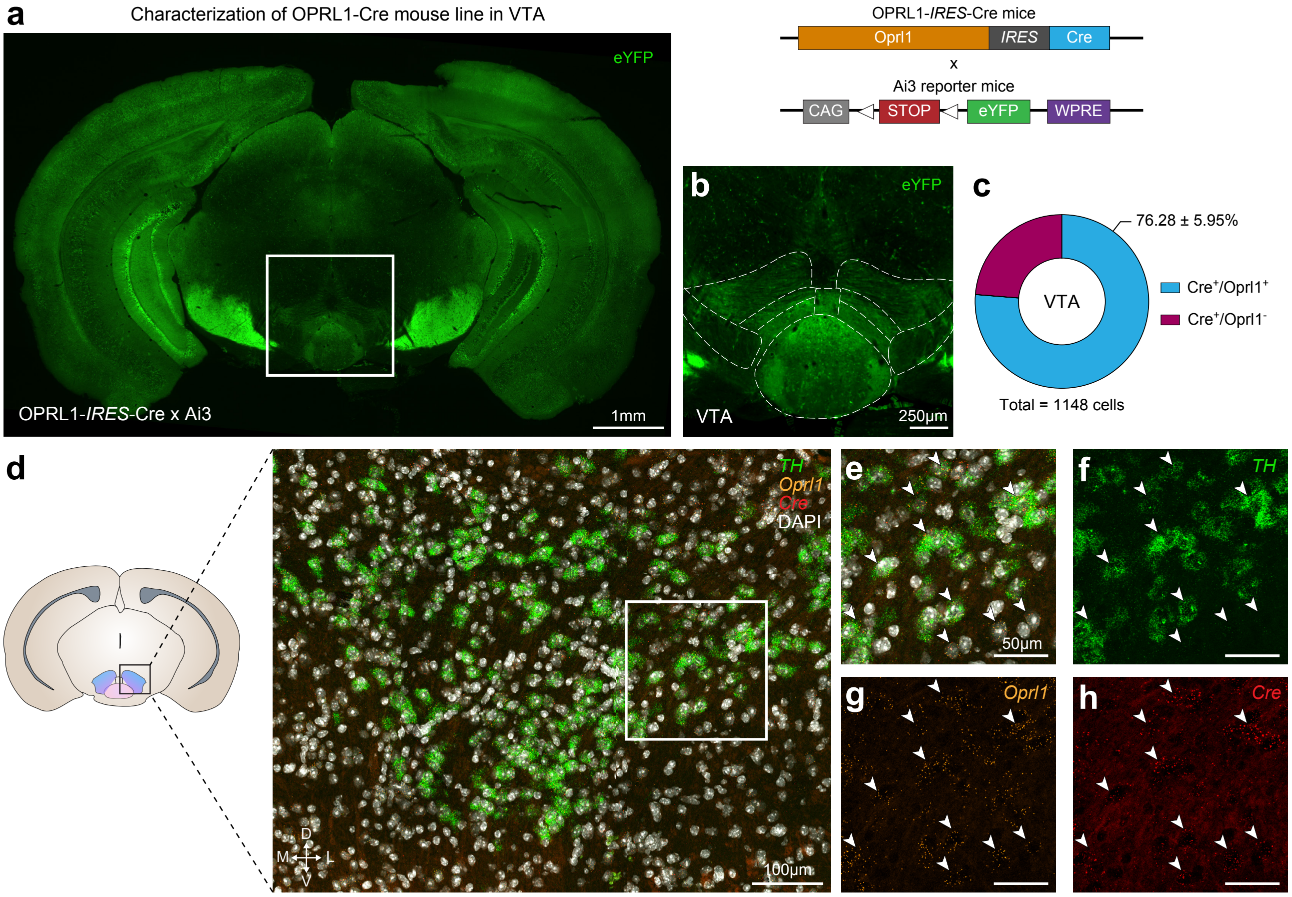


**SUPPLEMENTARY FIGURE 9 Anatomical characterization of the OPRL1-Cre mouse line in the ventral tegmental area.**
**a.** An OPRL1-Cre mouse line was generated and crossed with an Ai3 reporter line. Coronal image at bregma -3.28 mm showing Ai3-eYFP expression driven by OPRL1-Cre. **b.** Higher magnification of inset from **a** showing eYFP reporter expression in the ventral tegmental area (VTA). **c.** Quantification of *Oprl1* expression within Cre-expressing cells detected by *in situ* hybridization in the VTA (n = 2; 2 mice, 2-3 slices each). Data represented as mean ± SEM. **d.** Representative image showing *Oprl1* and Cre expression patterns in Oprl1-Cre mice via *in situ* hybridization of *Oprl1* (orange), Cre (red), *Th* (tyrosine hydroxylase, green), and DAPI (white) in the VTA. **e-h.** Higher magnification of inset from **d** showing combined (**e**) and individual (**f-h**) channels for *Th*, *Oprl1*, Cre, and DAPI. Scale bars, 50 μm.

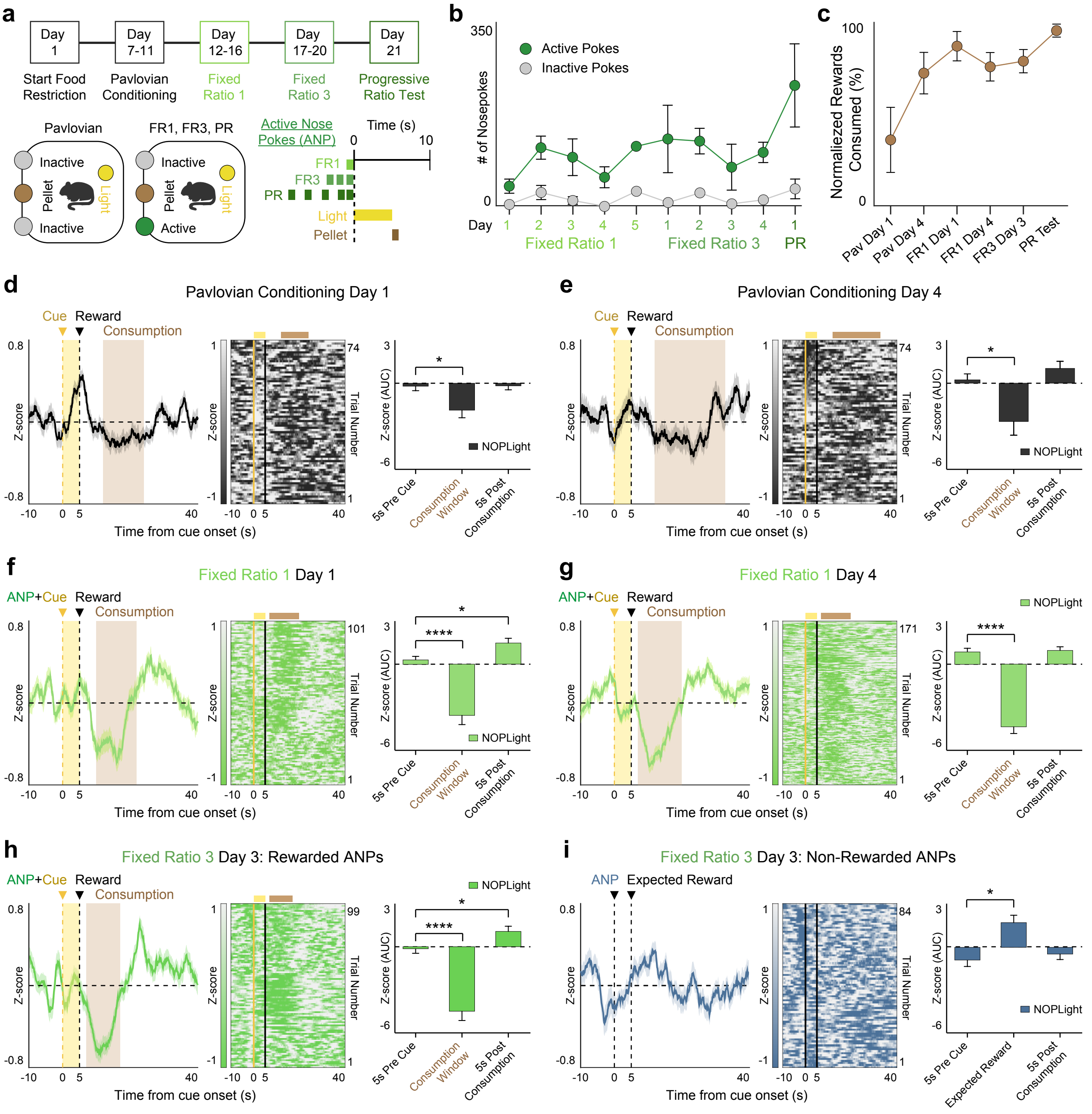


**SUPPLEMENTARY FIGURE 10** NOPLight detection of endogenous VTA N/OFQ release during Pavlovian and operant conditioning.

**SUPPLEMENTARY FIGURE 10 NOPLight detection of endogenous VTA N/OFQ release during Pavlovian and operant conditioning.** **a.** Mice expressing NOPLight in the VTA from Figure 6 were trained on Pavlovian and fixed ratio schedules prior to entering the progressive ratio test (n = 5 mice). Top: Timeline depicting training regimen for Pavlovian conditioning and operant conditioning. Bottom: Cartoon depicting operant box setup and the trial structure for each training paradigm. **b.** Total number of nosepokes made in the active (green) or inactive (gray) nosepoke ports across the entire training regimen. Data represented as mean ± SEM. **c.** Proportion of delivered rewards that were consumed by the mice on each of the photometry recording days. Data represented as mean ± SEM. **d-e.** Left: Trace of mean NOPLight signal during the reward period of the first (**d**) and fourth (**e**) Pavlovian conditioning sessions, aligned to start of light cue (yellow, shaded). Time to pellet retrieval and duration of consumption period averaged across all trials and animals (brown, shaded). Middle: Corresponding heat map. Right: Area under the curve (AUC) for photometry traces from **d** and **e** respectively, calculated over the averaged reward consumption time window and in 5-second intervals before/after the reward period (two-tailed Wilcoxon test, *$p<0.05$, n = 2-5 mice). Data represented as mean ± SEM. **f-g.** Left: Trace of mean NOPLight signal during the reward period of the first (**f**) and fourth (**g**) fixed ratio 1 (FR1) sessions, aligned to active nose pokes. Time to pellet retrieval and duration of consumption period averaged across all trials and animals (brown, shaded). Middle: Corresponding heat map. Right: Area under the curve (AUC) for photometry traces from **f** and **g** respectively, calculated over the averaged reward consumption time window and in 5-second intervals before/after the reward period (two-tailed Wilcoxon test, *$p<0.05$, **** $p<0.0001$, n = 3 mice). Data represented as mean ± SEM. **h.** Left: Trace of mean NOPLight signal during the reward period of a single fixed ratio 3 (FR3) session, aligned to reinforced active nose poke epoch. Middle: Corresponding heat map, where each row corresponds to an individual FR3 reward period. Time to pellet retrieval and duration of consumption period averaged across all trials and animals (brown, shaded). Right: Area under the curve (AUC) for the photometry trace, calculated over the averaged reward consumption time window and in 5-second intervals before/after the reward period (two-tailed Wilcoxon test, *$p<0.05$, **** $p<0.0001$, n = 3 mice). Data represented as mean ± SEM. **i.** Left: Trace of mean NOPLight signal during the same fixed ratio 3 (FR3) session as **h**, aligned to non-rewarded active nose pokes. Middle: Corresponding heat map, where each row corresponds to an individual, non-reinforced active nose poke epoch. Right: Area under the curve (AUC) for the photometry trace, calculated over the expected reward period, and in 5-second intervals before/after the expected reward period (two-tailed Wilcoxon test, *$p<0.05$, n = 3 mice). Data represented as mean ± SEM.
